# A Brighter picALuc Generated Through the Loss of a Salt Bridge Interaction

**DOI:** 10.1101/2023.02.14.528398

**Authors:** Kabir H Biswas

**Affiliations:** Division of Biological and Biomedical Sciences, College of Health & Life Sciences, Hamad Bin Khalifa University, Qatar Foundation, Doha – 34110, Qatar

**Keywords:** ALuc, Bioluminescence, Luciferase, Molecular Dynamics Simulation, picALuc

## Abstract

Recently, a miniaturized variant of an artificial luciferase (ALuc), named picALuc, with a molecular weight of 13 kDa and thus, the smallest luciferase, was reported. While picALuc was found to be as active as the ALuc, questions remained on the structural organization and residue-residue interactions in the protein. Here, combining structural modeling, molecular dynamics (MD) simulations and mutational analysis, we show that the loss of a salt bridge interaction formed by Glu50 (E50) residue results in an increased enzymatic activity of picALuc. Specifically, we generated a model of picALuc using the available structure of the *Gaussia* luciferase (GLuc) and performed a 1 μs long Gaussian accelerated molecular dynamics (GaMD) simulation which revealed a general compaction of the protein structure as well as residue level interactions in the protein. Given that picALuc contains a number of charged residues, we focused our attention to salt bridge interactions and decided to mutate E10, E50 and D94 that were found to form a fluctuating, stable or a new salt bridge interaction, respectively. Live cell assays showed an enhanced bioluminescence in cells expressing the E50A mutant picALuc while in vitro assays revealed an increased *V*_max_ of the E50A mutant without affecting its thermal stability. Dynamic cross-correlation and principal component analyses of the GaMD simulation trajectories revealed altered collective dynamics in the protein, in which residue E50 contributed substantially. Finally, we developed a protein fragment complementation assay using picALuc that allows monitoring protein-protein interaction in live cells. We envisage that the brighter variant of picALuc and the protein fragment complementation assay reported here will find a general applicability in developing bioluminescence-based assays and the strategy developed here will pave the way for further engineering of brighter variants of picALuc.

## Introduction

Bioluminescence has been utilized extensively in developing a wide range of biological applications including monitoring gene expression, protein-protein interaction, and protein structural changes.^2, 3, 4, 5, 6, 7, 8, 9^ These applications are primarily based on the activity of luciferase proteins that emit light at a characteristic wavelength upon oxidation of a cognate substrate, with or without requiring ATP. To further increase their utility in biological and biomedical applications, several luciferase variants have been characterized or developed with a range of spectral properties i.e. the peak emission wavelength. For example, mutational analysis, in combination with the use of different substrates, has enabled the generation of *Renilla* luciferase (RLuc) variant with a range of emission wavelengths including those that show peak emission in the red region of the spectrum.^10, 11, 12, 13, 14, 15^ Additionally, significant efforts have been made towards developing luciferases that show enhanced bioluminescence activity. These include variants that show an increased bioluminescence such as the consensus-based analysis, protein-protein interaction analysis and protease activity sensors^21, 23, 24^, including its application in protein fragment complementation assay for monitoring protein-protein interaction^25, 26, 27^

More recently, efforts were made to generate artificial luciferases (ALucs) using consensus amino acid sequence of copepod luciferases.^28^ This resulted in the generation of a number of ALucs with unique sequence and spectral properties such enhanced and stable bioluminescence. The ALucs were then utilized for the development of assays such as mammalian two-hybrid assay, live cell imaging and bioluminescent antibodies. Additional efforts towards generating ALucs using the same approach resulted in the generation of brighter variants with a preference for native coelenterazine as a substrate and some with alter spectral properties.^29^ Additionally, analysis of amino acid sequences of the ALucs revealed role for the C-terminal residues in bioluminescence activity of the ALucs but not in their spectral properties. The latest addition in this series of luciferases is a miniaturized version of one of the ALucs (ALuc30) mutational analysis of RLuc led to the development of Rluc8^16^ and Super Rluc8^17^. More recently, insertion and deletion mutagenesis of an ancestral RLuc allowed generation of an RLuc variant with a highly stable, glow-type bioluminescence from an otherwise characteristic flash-type bioluminescence.^18^

In the recent past, generation of NanoLuc (NLuc) from the 19 kDa subunit of the deep-sea shrimp (*Oplophorus gracilirostris*) luciferase has been a major success due to its smaller size, higher and prolonged bioluminescence activity and greater thermal stability.^19, 20, 21, 22^ These advantageous features of the enzyme have led to its utilization in a range of applications including gene expression generated previously.^30^ Aptly referred to as picALuc due to its size (smallest among luciferases so far with a molecular weight of 13 kDa), it was generated through the deletion of residues 1-54 at the N-terminal and residues 175-194 at the C-terminal of ALuc30. Importantly, picALuc was reported to be as active as the parental ALuc30 and showed thermal stability and brightness similar to that of NLuc, thus raising the possibility of its utilization in all bioluminescence-based assays including Bioluminescence Resonance Energy Transfer (BRET).^30, 31, 32, 33, 34^

In the current study, we have generated a structural model of picALuc using the structure of GLuc^1^ as a template, which revealed a large ‘hole’ in the N-terminal part of the protein due to the absence of helices *α*1 and *α*2 that were present in the template structure. We then performed an all-atom, explicit solvent, 1 *μs*-long Gaussian accelerated MD (GaMD) simulations to understand structural features and inter-residue interactions in the protein. The simulation revealed structural compaction of the protein early on during the simulation, largely due to the movement of the N-terminal *α* helices of picALuc into the positions occupied by *α* helices deleted in picALuc. Importantly, we observed increased number of salt bridge interactions in the protein and mutation of salt bridge interaction forming residues E10, E50 and D94 to A revealed a large increase in the bioluminescence activity of picALuc with the E50A mutation. Biochemical and biophysical characterization revealed that the increased bioluminescence activity of the E50A mutant is due an increase in the *V*_max_ of the protein without any significant alteration in the thermal stability of the protein. We then performed MD simulation of the E50A mutant picALuc compared it with that of the WT protein revealing differences in the collective dynamics in the protein, in addition to much reduced contribution of the A50 residue in the E50A mutant picALuc compared to E50 in the WT protein. Finally, we report the development of a picALuc-based protein fragment complementation assay that allows monitoring protein-protein interaction in live cells.

## Results & Discussions

### Molecular dynamics simulation reveals picALuc structural features and reorganization

Deletion mutagenesis of ALuc, a synthetically designed luciferase protein, resulted in the generation of the smallest, 13 kDa, luciferase, picALuc^30^. Specifically, the authors deleted residues 1 to 54 at the N-terminal (comprising of helices *α*1 and *α*2) and residues 175 to 194 at the C-terminal side (comprising of an extended unstructured loop region) (Supplementary Figure 1B). To better understand the structural features of picALuc, we generated a structural model of the protein using the available NMR structure of GLuc. For this, the amino acid sequences of the picALuc and GLuc were aligned (Supplementary Figure 1A) and a structural model was generated using the Modeller software (Fig. 1A-D). One of the key features of GLuc is the presence of five disulfide bridges formed by residue pairs C28-C95, C31-C92, C37-C49, C24-C99 and C108-C120 (Fig. 1A and Supplementary Figure 1C), which were maintained in the picALuc model. Additionally, all secondary structural elements were maintained in the modelled structure of picALuc (Fig. 1A, B), except for those which were deleted in picALuc (*α* helices in Fig. 1B were numbered according to the picALuc sequence). Overall, GLuc and the modelled picALuc structures showed a C*α* RMSD value of 1.5 Å (Fig. 1C). However, a closer inspection of the two structures revealed a non-compact structure with a ‘hole’ at the N-terminal side due to the absence of the helices *α*1 and *α*2, which are present in the GLuc structure but deleted in picALuc (Fig. 1D).

**Fig.1.**
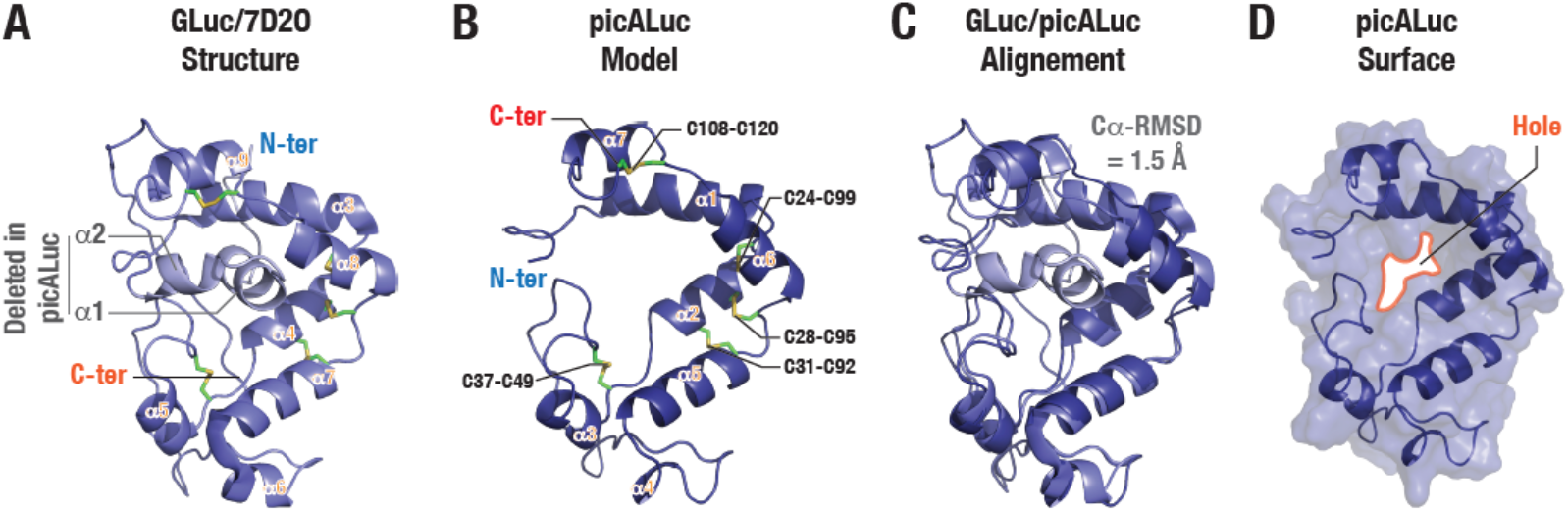
picALuc sequence and structural features. (A) Cartoon representation of GLuc structure highlighting various secondary structural elements (α-helices), disulfide bridges and the equivalent N-terminal region deleted in picALuc. (B) Cartoon representation of picALuc structural model generated using the available GLuc structure (PDB: 7D2O)^1^. Disulfide bridges, secondary structural elements (α-helices) and N and C-termini. (C) Cartoon representation of aligned GLuc and modelled picALuc structures. (D) Cartoon and surface representation of the modelled picALuc structure showing the presence of a ‘hole’ due to the absence of the N-terminal region (helices α1 and α2 of GLuc).

To determine if the modelled picALuc structure is stable or assumes a more compact form, we performed molecular dynamics (MD) simulations. For this, we set up an all atom, explicit solvent MD simulation and ran the simulation for 1 μs using NAMD^35^. Importantly, we employed GaMD simulation to better explore the conformational dynamics of the protein within the simulation period of 1 μs (Supplementary Movie 1).^36, 37^ Analysis of the simulation trajectory revealed a rapid compaction of the picALuc structure within the first 10 ns of the simulation with further changes observed until 50 ns (Fig. 2A). Root mean square distance (RMSD) of C*α* atoms revealed large changes in the initial phases of the simulation with stabilization of the RMSD trace observed after about 100 ns. Additionally, root mean square fluctuation (RMSF) analysis of the trajectory revealed largest fluctuations in residues corresponding to the loop between helices *α*3 and *α*4 while residues corresponding to helix *α*3 in the N-terminal side and helices *α*6 and *α*7 in the C-terminal side (Fig. 2C). Further, pairwise RMSD analysis of the trajectory also captured the large structural changes in picALuc in the initial 100 ns of the simulation with largest changes seen in the first 10 ns of the simulation (Fig. 2D).

**Fig. 2.**
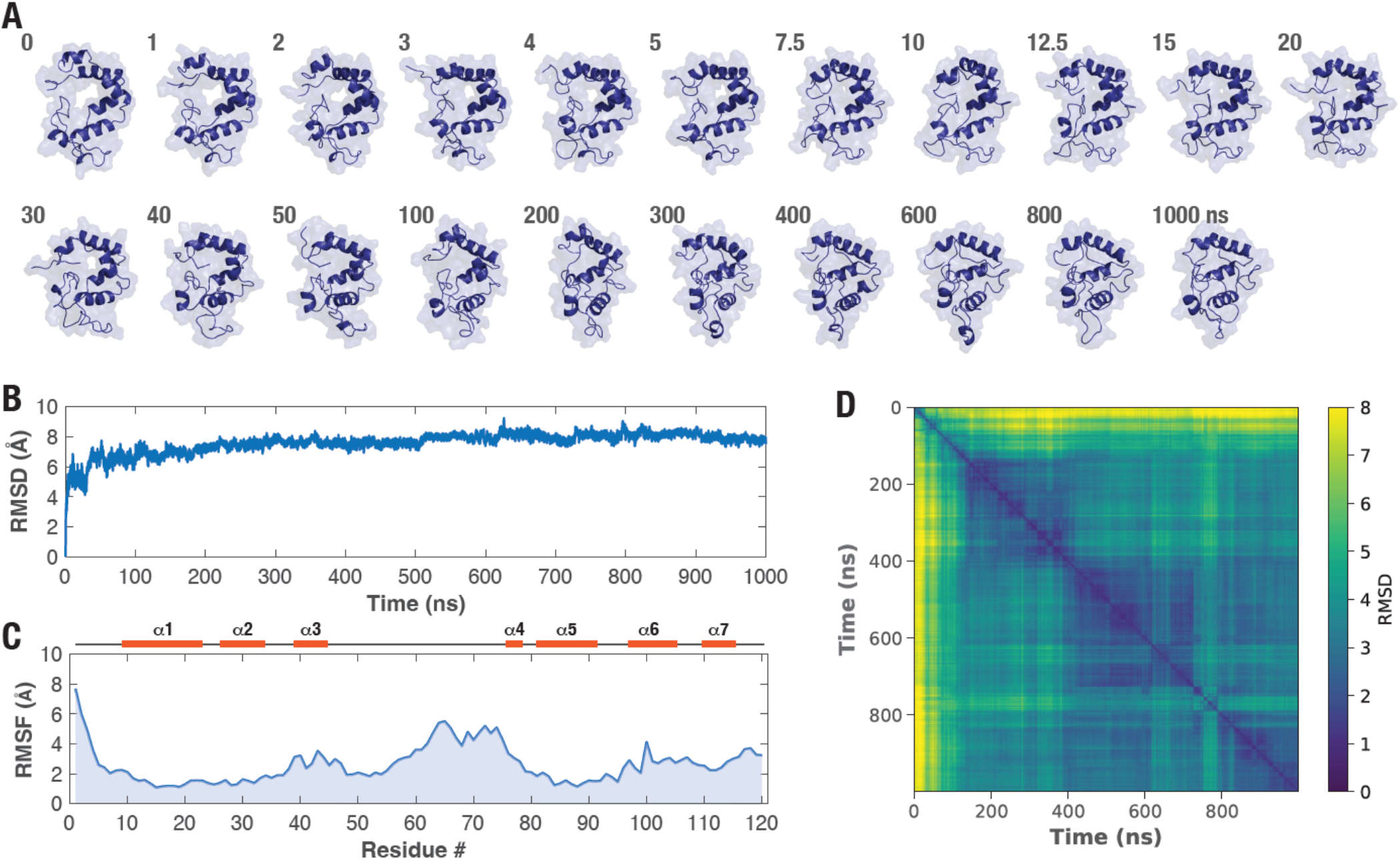
MD simulation reveals structural reorganization in picALuc. (A) Cartoon and surface representation of picALuc at the indicated period over the course of 1 μs GaMD simulation showing rapid structural evolution of the protein. Note that the frames shown are to best capture the structural during the initial phase of the simulation and the time difference is not uniform. (B,C) Graphs showing Cα atom RMSD (B) and RMSF (C) values of picALuc obtained from the GaMD simulation. Secondary structural elements (α helices) are highlighted in the RMSD graph for reference. (D) Heat map showing pairwise RMSD of Cα atom of picALuc. Note significant changes in the initial phases of the simulation. RMSD values are in Å.

We then performed solvent accessible surface area (SASA) and radius of gyration (RoG) analysis of the protein to confirm the structural compaction. These revealed a general reduction in both SASA and RoG during the initial 200 ns of the simulation with somewhat slower decrease in the initial 100 ns but a much rapid decrease in between 100 and 200 ns (Fig. 3A,B). This is contrast with the C*α* RMSD and pairwise RMSD analyses which showed large changes during the initial 100 ns and likely reflects side chain rearrangements and further packing in the protein. We also determined various types of energies of the protein (Fig. 3C,D; Supplementary Figure 3) and found that the van der Waal energy decreased in the initial phases of the simulation but reverted back to similar level during the later phases of the simulation. On the other, electrostatic (as well as non-bond and total) energy of the protein decreased during the initial phases of the simulation and remained low during the later phases of the simulation. However, the number of hydrogen bonds (H-bonds) also remained similar during the simulation (Fig. 3E; Supplementary Table 1). Additionally, secondary structure analysis revealed changes only in the C-terminal part of the loop between helices *α*3 and *α*4 (Fig. 3F; Supplementary Figure 4), indicating that the structural compaction of picALuc observed during the simulation is likely due to movement of the initially modelled *α* helices. We note that all five disulfide bridges were maintained all throughout the simulation (Supplementary Figure 5).

**Fig. 3.**
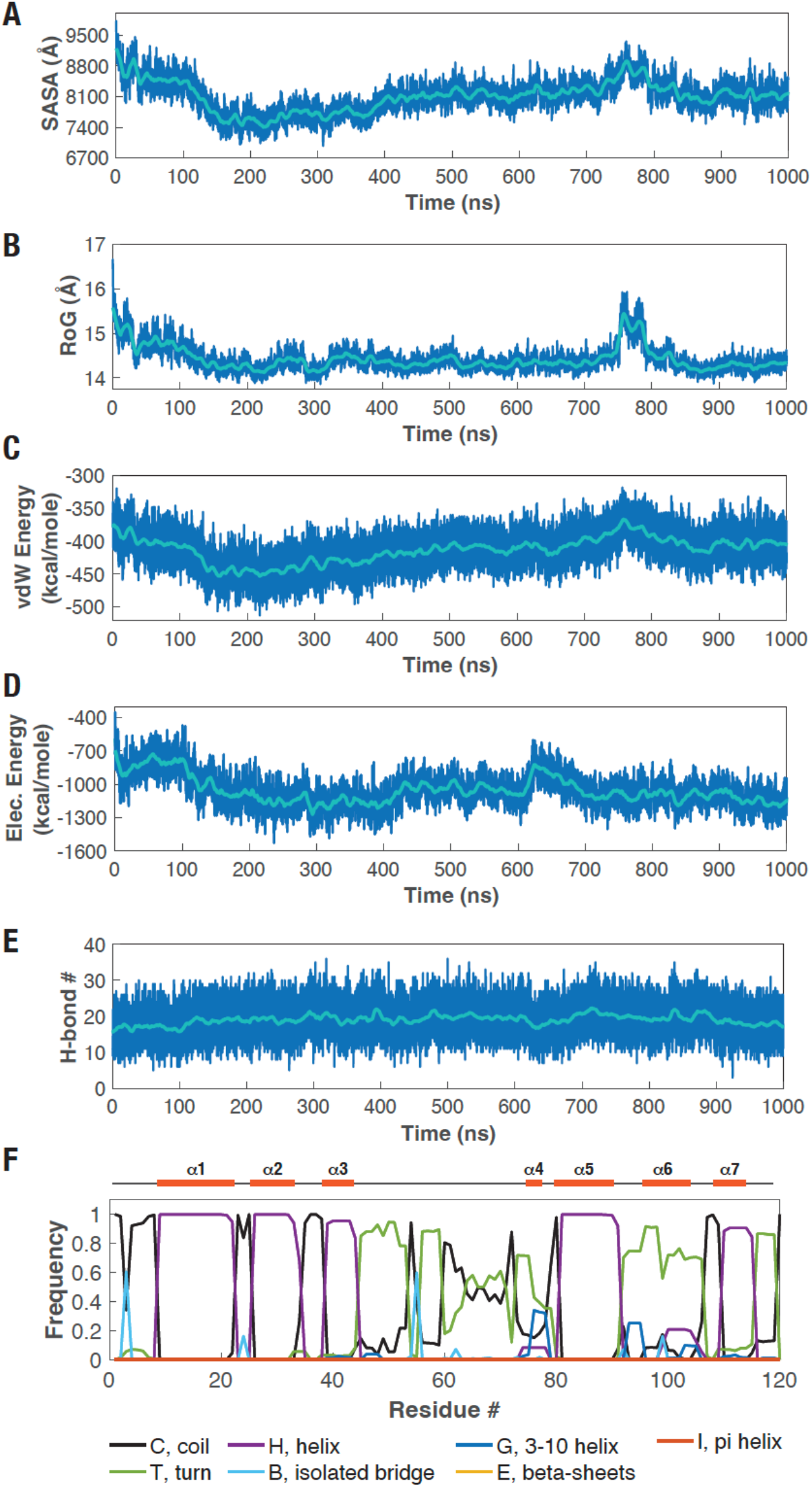
Structural and energetic features of picALuc. (A,D) Graphs showing solvent accessible surface area (SASA) (A), radius of gyration (RoG) (B), van der Waal’s energy (vdW Energy) (C) and electrostatic energy (D) of picALuc determined from a 1 μs-long GaMD simulation. Note the general reduction in each of these parameters during the initial phases of the simulation (0-200 ns). (E) Graph showing number of H-bonds formed over the course of 1 μs GaMD simulation. (F) Graph showing the frequency of various secondary structural elements in picALuc over the course of 1 μs GaMD simulation.

### Increased number of salt bridge interactions formed in picALuc

Given the observation of increased electrostatic interaction (Fig. 3D) and the presence of a large number of charged residues in picALuc (Supplementary Figure 1B), we then focused our attention on the salt bridge interactions formed during the 1 *μs* long GaMD simulation. Determining the total number of salt bridge interactions formed with an oxygen-nitrogen distance cutoff of 3.2 Å during the simulation revealed an increase in the number of salt bridge interactions formed by the protein (Fig. 4A), with largest increases observed during the initial phase of the simulation. We, therefore, analyzed the trajectory for salt bridge interactions formed during the simulation and found several salt bridge interactions that were found during the initial phase of the simulation while a number of such interactions were lost (Supplementary Figure 6). In addition, several salt bridge interactions were found to be fluctuating or were formed transiently (Supplementary Figure 7). For instance, residues E10 and K13 (both located in the helix *α*1) formed a salt bridge interaction that showed fluctuating distance between the two amino acid residues with a mean (± s.d.) and median distance of 8.69 ± 3.64 and 10.01 Å, respectively and a 16% fractional occupancy below 3.2 Å (Fig. 4B). Importantly, we observed relatively stable salt bridge interactions formed between E16 and R26, E50 and K36 and E50 and K42 throughout the trajectory. For instance, residues E50 (located in the loop between helices *α*3 and *α*4) and K42 (located in the helix *α*3) formed a relatively stable salt bridge interaction with a mean (± s.d.) and median distance of 4.57 ± 1.96 and 3.58 Å, respectively and a 23% fractional occupancy below 3.2 Å (Fig. 4C). On the other hand, residues D94 (located in the loop between helices *α*5 and *α*6) and K56 (located in the loop between helices *α*3 and *α*4) formed a new salt bridge interaction during the initial phase of the simulation with a mean (± s.d.) and median distances of 4.75 ± 4.86 and 3.36 Å, respectively and a 32% fractional occupancy below 3.2 Å (Fig. 4D).

**Fig. 4.**
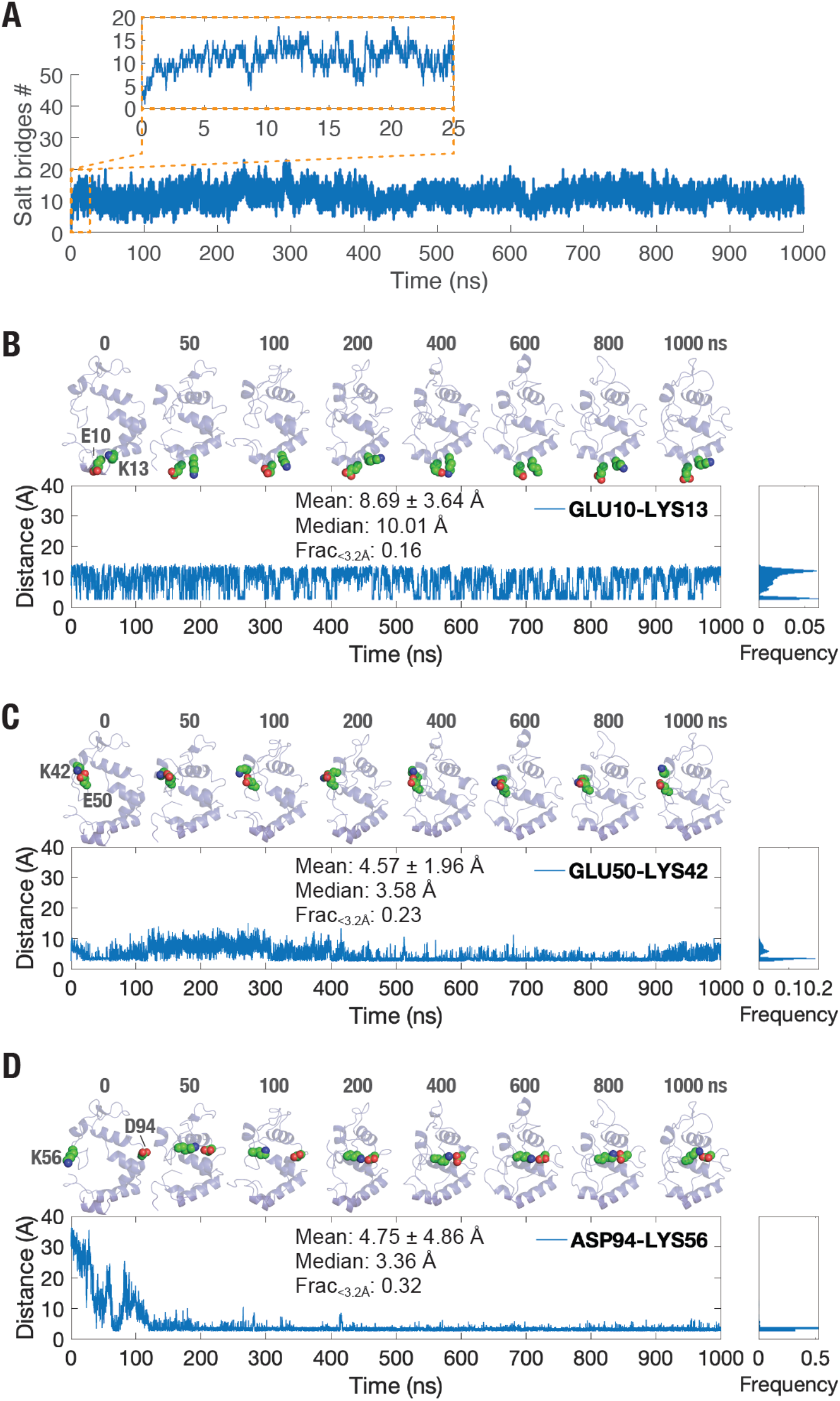
Salt bridge interactions in picALuc. (A) Graph showing number of salt bridges formed during a 1 μs long GaMD simulation. Inset shows increase in the number of salt bridges during the early phases of the simulation (0-25 ns). (B-D) Cartoon representation (top panels) and graphs showing inter-residue distances (bottom panels) of salt bridge forming residue pairs Glu10 and Lys13 (B), Glu50 and Lys42 (C) and Asp94 and Lys56 (D). Mean and median distances and fractional occupancies of each of the salt bridges are shown as insets. Note that the picALuc structure has been reoriented to best show the salt bridge interactions.

### Mutational analysis reveals increased bioluminescence of picALuc

Following our observations with the specific salt bridge interactions above, we decided to determine their role in the bioluminescence activity of picALuc. For this, we mutated the residues E10, E50 and D94 to A (E10A, E50A and D94A, respectively) and expressed the proteins, along with the wild type (WT) protein, in mammalian (HEK 293T) cells. Additionally, we fused the mGreenLantern (mGL) green fluorescent protein at the N-terminal of picALuc for monitoring expression level of the proteins (Fig. 5A). Fluorescence measurement of living cells expressing the WT and the mutant picALuc proteins showed similar expression of the WT, E10A and E50A mutants while the D94A mutant showed higher levels of the protein (Fig. 5B). Bioluminescence measurement of the cells, on the other hand, showed similar activity of the WT and E10A and D94A mutant picALuc while the E50A mutant showed higher activity (Fig. 5C). Bioluminescence spectral measurements of the cells expressing the proteins showed a single peak at around 526 nm (Fig. 5D) likely indicating a high efficiency of resonance energy transfer measured as a ratio of bioluminescence (Fig. 5E). Indeed, inclusion of the self-cleaving P2A peptide^38, 39^ in between mGL and picALuc resulted in a blue-shifted spectra and a decrease in BRET (Supplementary Figure 8). Additionally, proteolytic cleavage of the SARS-CoV-2 M^pro^ N-terminal autocleavage site present in between mGL and picALuc upon co-expression of the M^pro^ resulted in shift in the spectra and a significant decrease in the BRET (Supplementary Figure 9).^31, 32, 33, 40, 41, 42^ Overall, we observed that the E50A mutant picALuc showed about 3.8 times bioluminescence activity (after normalization with total protein levels as determined from mGL fluorescence) compared to the WT picALuc (Fig. 5F). Thus, while the residue E50 formed a stable salt bridge interaction (with the residue K42) throughout the 1 *μs* GaMD simulation, mutation of the same resulted in an increased bioluminescence activity of picALuc.

**Fig. 5.**
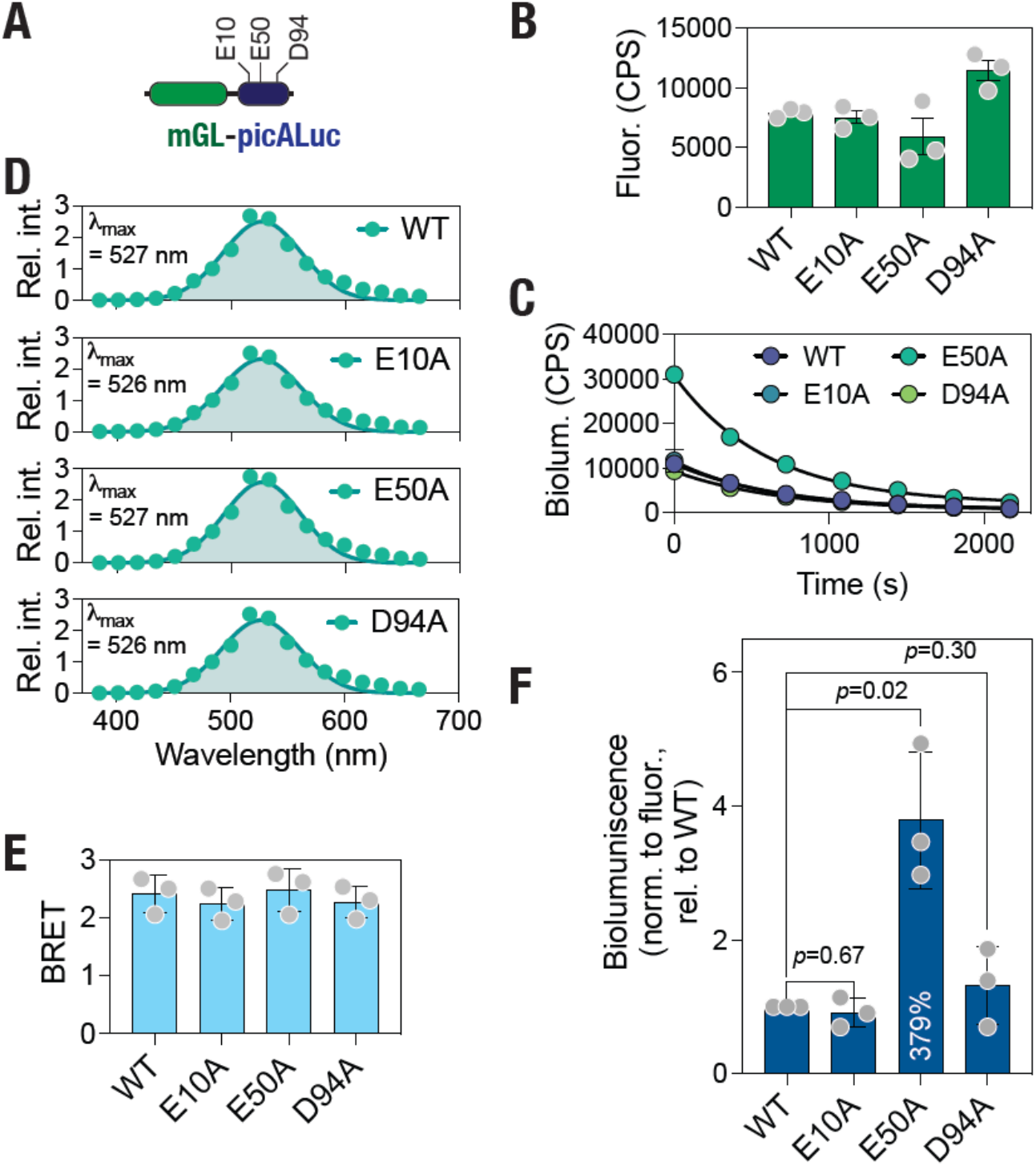
Increased bioluminescence of the E50A mutant picALuc. (A) Schematic showing the organization of the mGL-picALuc protein highlighting the relative location of E10, E50 and D94 residues in the protein. (B,C) Graphs showing fluorescence (B) and bioluminescence (C) in cells expressing either the WT or mutant mGL-picALuc proteins. Data shown are mean ± s.e.m of measurements from a representative experiment performed in triplicates. (D) Graphs showing bioluminescence spectra of cells expressing either the WT and mutant mGL-picALuc proteins. Bioluminescence data were fit to a Gaussian distribution. (E) Graph showing BRET values of cells expressing either the WT or the mutant mGL-picALuc proteins. (F) Graph showing bioluminescence (normalized to fluorescence values and relative to the WT mGL-picALuc) of cells expressing either the WT or the mutant mGL-picALuc proteins. Data shown in E and F are mean ± s.d. from three independent experiments performed in triplicates.

### E50A mutation results an increased enzymatic activity without altering thermal stability of the protein

Having observed an increased bioluminescence activity of the E50A mutant picALuc in living cells, we decided to determine the biochemical and thermal property of the protein in vitro. For this, HEK 293T cells transfected with the WT and various mutant picALuc proteins were used for preparing picALuc containing cell lysates to be used for in vitro assays. First, we performed enzyme kinetic studies with the WT and various mutants (equivalent concentrations of proteins were used) under a range of substrate (coelenterazine h) concentrations and utilized rate of photon emission as the enzymatic rate (bioluminescence; counts per second, CPS) and fitted the data to an allosteric sigmoidal model (Fig. 6A). Consistent with the results obtained with live cells, we observed a higher bioluminescence activity with the E50A mutant picALuc compared to the WT picALuc. While the rate constant (*K*_half_) was found to be similar for all four proteins (Fig. 6B), the maximum enzyme velocity (*V*_max_) was significantly higher for the E50A mutant compared to the WT picALuc (Fig. 6C). Additionally, we observed a significant decrease in the *V*_max_ of the E10A and D94A mutant compared to the WT picALuc (Fig. 6C), suggesting a role for these residues and likely the salt bridge interactions formed by these residues in the catalytic activity of picALuc.

**Fig. 6.**
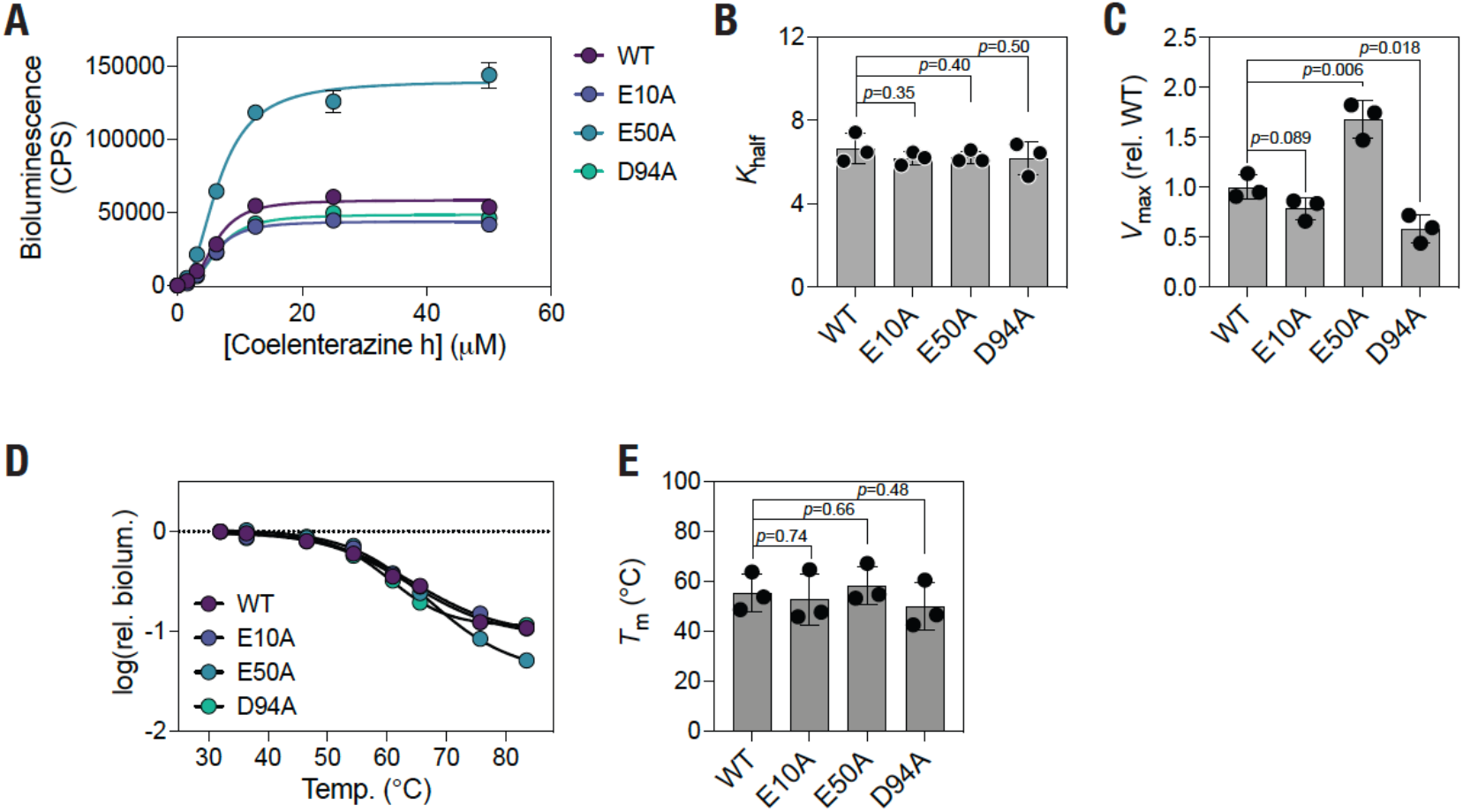
E50A mutation increases picALuc V_max_ without altering thermal stability. (A) Graph showing bioluminescence activity in lysates prepared from cells expressing either the WT or mutant mGL-picALuc proteins under the indicated substrate (coelenterazine h) concentrations. Data shown are mean ± s.e.m of measurements from a representative experiment and fit to an allosteric sigmoidal model. (B,C) Graphs showing K_half_ and V_max_ values of the WT and mutant mGL-picALuc proteins. Data shown are mean ± s.d. from three independent experiments. (D) Graph showing relative bioluminescence activity of either the WT or the mutant mGL-picALuc protein incubated at the indicated temperatures. (E) Graph showing melting temperature of WT and mutant picALuc proteins obtained from Boltzmann sigmoidal model fitting of melting temperature curves. Data shown are mean ± s.d. from three independent experiments.

Further, we determined the bioluminescence activity of the four picALuc proteins after incubation at a range of temperatures (from 32 to 84 °C) for 10 min. We fitted the data to a Boltzmann sigmoidal model to determine the melting temperature (*T*_m_) of the proteins (Fig. 6D). We observed a relatively high thermal stability of picALuc with a *T*_m_ of 55.4 ± 7.7 °C (Fig. 6E). However, no significant changes were observed in the *T*_m_ of the mutants, including E50A (Fig. 6E). This likely reflects the thermal stability provided by the five disulfide bridges in the protein. Taken together, these data suggest that the E50A mutation increases the bioluminescence activity of picALuc by increasing its *V*_max_ without affecting its thermal stability.

### Altered structural dynamics in the E50A mutant picALuc

The increased bioluminescence activity of the E50A mutant picALuc was intriguing and therefore, we attempted to understand the underlying mechanism. For this, we introduced the E50A mutation in the structural model of picALuc and performed an all atom, explicit solvent 1 *μs* GaMD simulation similar to the WT protein (Supplementary Movie 2). We then analyzed the trajectories of the WT and the E50A mutant picALuc to determine the distance between the C*α* atoms in residues at position 50 (E in the WT and A in the E50A mutant picALuc) and 42 (K in both proteins) and found that the E50A mutation did not result in any major changes between the two proteins except for some differences in the initial phases of the simulation (Fig. 7A), likely reflects the overall structural stability of the mutant protein. We then analyzed the E50A mutant picALuc trajectory for Cα RMSF values and mapped the RMSF values of individual amino acid residues onto the structural conformer obtained from the last frame of the simulation as b-factors for the both the E50A mutant and WT picALuc. This revealed an increased fluctuation in the N-terminal residues and the long loop between helices *α*3 and *α*4 in the E50A mutant picALuc compared to the WT (Fig. 7B). We then performed dynamic cross-correlation (DCC) analysis of residues using the MD-TASK suite of MD simulation analysis software^43^ to elucidate any alterations in the correlated motions in the WT and E50A mutant picALuc proteins.^44^ The WT picALuc showed both positively and negatively correlated motions in the residues with prominent correlations observed for residues close to the residue E50 (Fig. 7C). Importantly, however, such highly positively correlated motions were lost in the E50A mutant picALuc while some correlated motions were gained between the range of residues 36-49 and 6-17 (Fig. 7C), suggesting a role for the residue E50 in the dynamics of the protein.

**Fig. 7.**
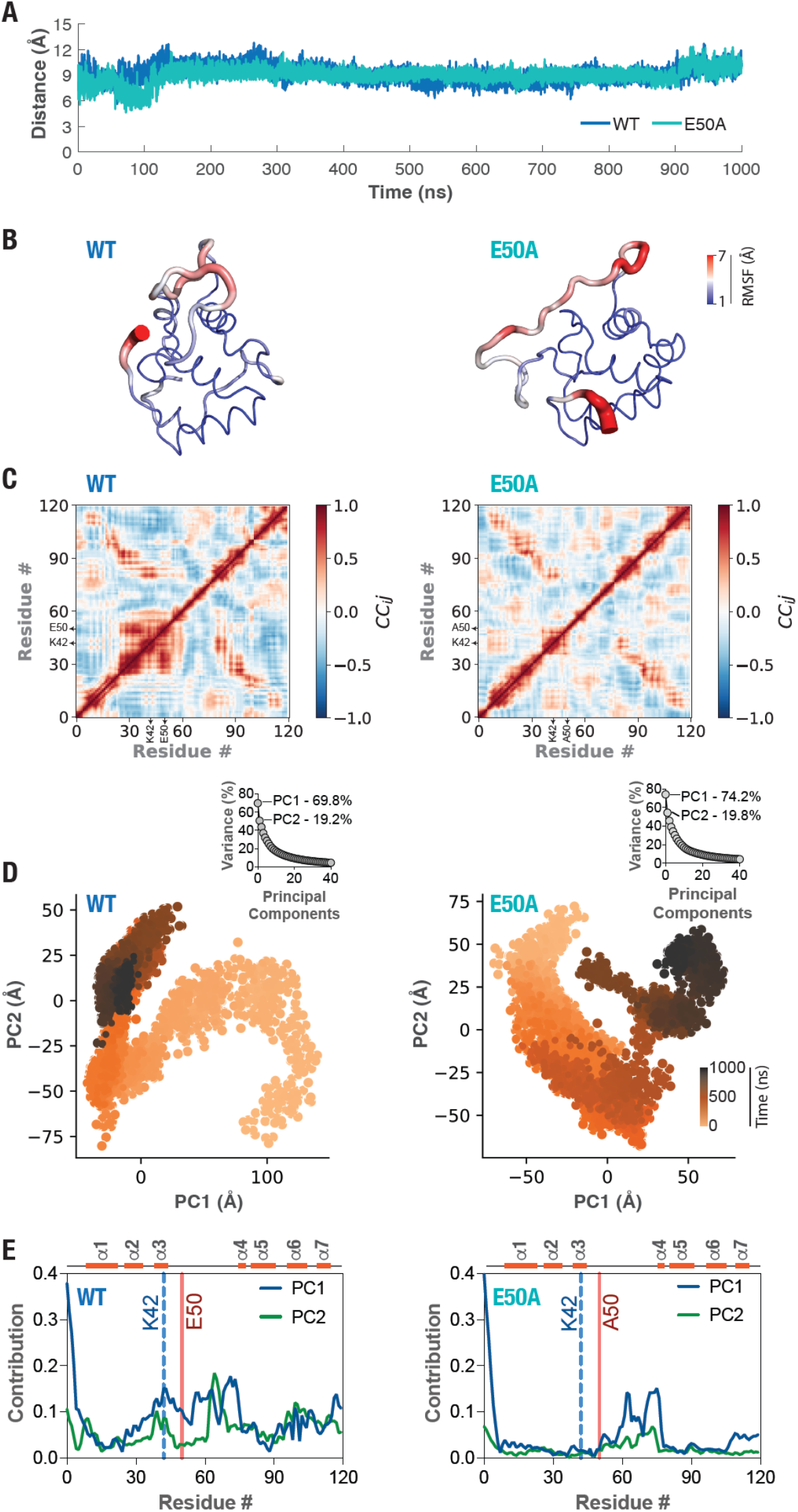
Altered structural dynamics in the E50A mutant picALuc. (A) Graph showing Cα atom distances between positions 50 and 42 (E50-K42 in the WT and A50-K42 in the E50A mutant) of picALuc over a 1 μs GaMD simulation. (B) Cartoon representation of the WT (left panel) and E50A mutant (right panel) picALuc with Cα atom RMSF values mapped on the conformers obtained after 1 μs of GaMD simulation (frame # 50,000) as b-factors. Color bar, Cα atom RMSF (Å). (C) Plots showing DCC of residues in the WT (left panel) and E50A mutant (right panel) picALuc determined from 1 μs GaMD trajectories of each protein. (D) Graphs showing principal components 1 and 2 (PC1 and PC2) of the WT (left panel) and E50A mutant (right panel) determined from 1 μs GaMD trajectories of each protein. Color bar, time (ns). Insets: graphs showing percentage variance against principal components obtained from PCA analysis of the WT (left panel) and the E50A mutant (right panel) picALuc. (E) Graphs showing contribution of individual residues to the PC1 and PC2 in the WT (left panel) and the E50A mutant (right panel) picALuc simulations. Locations of residue number 42 and 50 are highlighted using a blue and red line, respectively in each graph.

In order to further understand the structural dynamics of the WT and E50A mutant picALuc, we utilized the dimensionality reduction analysis and performed principal component analysis (PCA) of their trajectories, which is known to reveal collective motions in proteins^45^ using the PCA module available in the Python-based MDAnalysis package^46, 47^. Cumulative variance analysis of the principal components revealed the presence dominant collective motions in both the WT and the E50A mutant picALuc (Supplementary Figure 10) since principal components 1 and 2 (PC1 and PC2) could account for 69.8 and 19.2% of the variance in the WT protein while they could account for 74.2 and 19.8% of the variance in the E50A mutant protein. Importantly, this analysis revealed large differences in the PC1 and PC2 of the two trajectories (Fig. 7C), suggesting changes in the collective dynamics of picALuc upon E50A mutation. We, therefore, determined the contribution of individual residues to the PC1 and 2 in the WT and the E50A mutant picALuc trajectory and found residues K42 and E50, other than the N-terminal residues and those from the loop *α*3-*α*4, contributing significantly to PC1 in the WT protein (Fig. 7D). Importantly, such contributions from both residue positions (K42 and A50) as well as those from helix *α*3 in general were lost in the PC1 of the E50A mutant picALuc (Fig. 7D). These data suggest a role for the salt bridge interaction formed by residues E50 and K42 in the collective dynamics of picALuc, and that this mode of collective motion likely restricts the catalytic activity of the protein.

### Design of a picALuc-based split protein for monitoring protein-protein interaction in living cells

Finally, we attempted to design a split protein for protein fragment complementation-based assay for monitoring protein-protein interaction based on the picALuc conformer at 800 ns of the MD simulation (Fig. 2A). Specifically, we chose to utilize N-terminal residues spanning the helix *α*1 (Fig. 1B) until residue A22 (one G residue N-terminal to the first disulfide bond forming C24 residue) (Supp. Fig. 1C) as the smaller fragment named picSm (22 residue long; predicted molecular weight of 2.4 kDa) and the rest of the protein spanning residues G23 until C120 as the larger fragment named picLg (98 residue long; predicted molecular weight of 10.5 kDa) (Fig. 8A). For this, we designed a plasmid construct expressing the picSm fused with mGL at the N-terminal for monitoring expression levels of the protein and another expressing the picLg. Additionally, we generated a construct expressing mGL-picSm and picLg fused with the 19 residue long GCN4 leucine zipper peptide that forms a parallel, two-stranded coiled-coil structure^48, 49^ and has been utilized for synthetic dimerization applications including in BRET-based assays^25^. The latter two constructs were named mGL-picSm-GCN4 and picLg-GCN4, respectively. Transfection of HEK293T cells with increasing amounts of either mGL-picSm or mGL-picSm-GCN4 plasmid DNA, while maintaining the amount of picLg or picLg-GNC4, resulted in the increasing mGL fluorescence indicating increased expression of the picSm and picSm-GCN4 fragments (Fig. 8B). Importantly, transfection of cells with increasing concentration of mGL-picSm-GCN4 resulted in increased bioluminescence, while such increases were not observed in cells transfected with mGL-picSm (Fig. 8C). These results suggest that while picSm fragment likely do not possess high enough affinity for picLg to associate with it and drive bioluminescent activity of the complemented protein, GCN4-mediated dimerization likely brings the two fragments closer, enabling complementation and thus, bioluminescence activity. We note that while the picALuc SmBit is longer than the reported NanoLuc SmBit (22 vs. 11 residues), the picALuc LgBit is smaller than the NanoLuc SmBit (10.5 vs. 17.6 kDa)^50^, and thus may serve as a better fusion tag for monitoring protein-protein interaction, especially in the case proteins that are affected by fusion with larger protein tags.

**Fig. 8.**
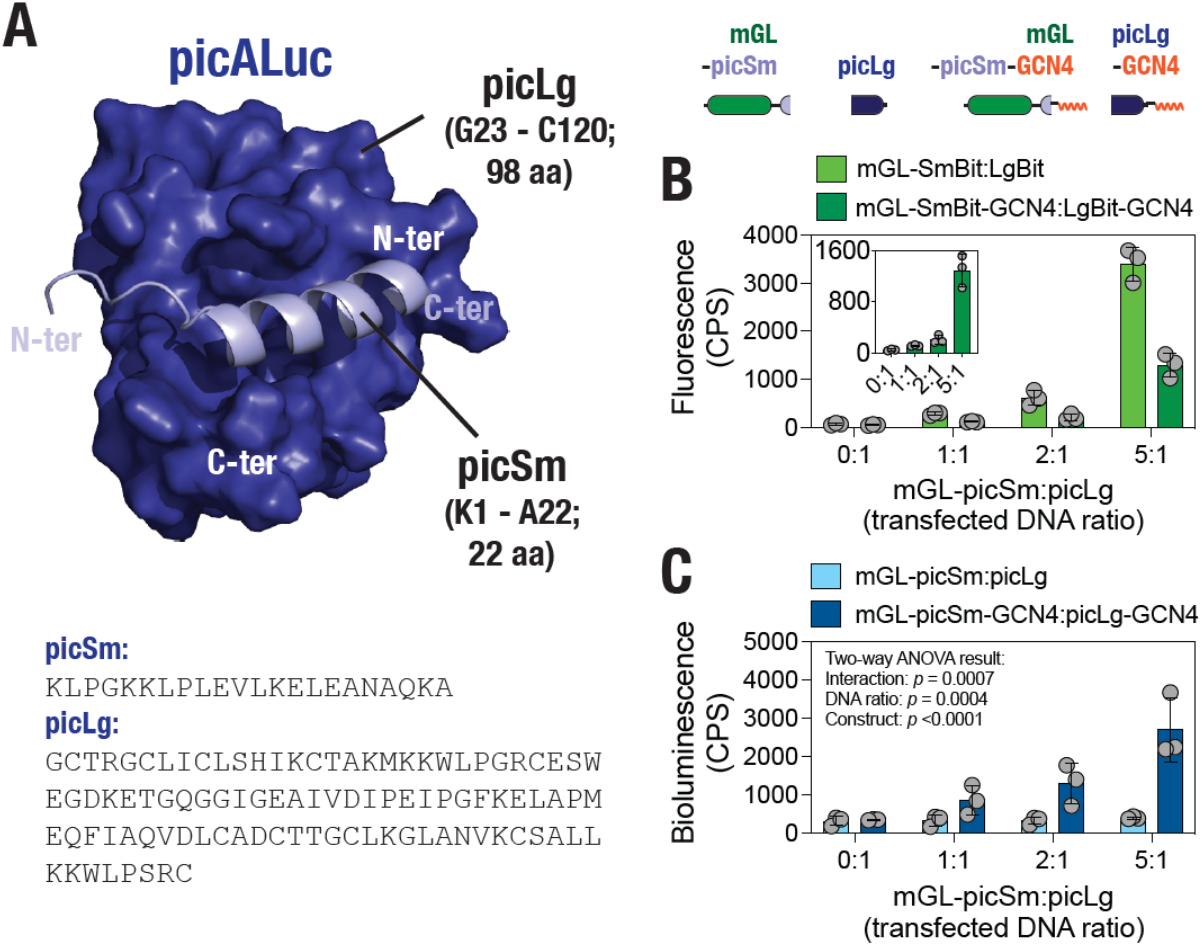
picALuc-based split protein for monitoring protein-protein interaction. (A) Cartoon representation of a picALuc conformer showing the N-terminal, α-helical, small (picSm) and large (picLg) fragments generated for monitoring protein-protein interaction. Bottom panel: Sequences of the picSm and picLg fragments. (B) Top panel: Schematic showing constructs generated for monitoring protein-protein interaction (dimerization) mediated bioluminescence activity. Bottom panel: Graph showing fluorescence in cells transfected with the indicated ratios of picSM and picLg DNA. Inset: fluorescence in cells transfected with picSm-GCN4 and picLg-GCN4. Data shown are mean ± s.d. of 3 measurements. (C) Graph showing bioluminescence activity in cells transfected with either mGL-picSm and picLg or mGL-picSm-GCN4 and picLg-GCN4 plasmid DNA at the indicated ratio (amount of plasmid DNA) showing increase in bioluminescence with increasing amount of the GCN4-fused picALuc fragments. Data shown are mean ± s.d. from 3 independent experiments, with each experiment performed in triplicates. Two-way ANOVA values show that changes in bioluminescence activity with increasing DNA ratios as well as between picSm:picLg and picSm-GCN4:picLg-GCN4 are significantly different.

## Conclusion

To conclude, MD simulation followed by detailed analysis of the trajectory not only revealed a compact three-dimensional structure of picALuc but also provided insights into the possible residue-level interactions. Specifically, closer inspection and mutational analysis of selected salt bridge forming residues enabled the generation of a brighter picALuc variant. Comparative analysis of the WT and E50A mutant picALuc MD simulation trajectories revealed an altered collective motion and specifically the contribution of positions 42 and 50 in the mutant picALuc. Finally, structural analysis allowed us to design a picALuc-based protein fragmentation assay that will find utility in monitoring protein-protein interaction. We believe that the results present here will pave the way for the generation of new variants of the smallest luciferase proteins. This will certainly increase the applicability of the luciferase in assays ranging from gene expression, protein stability, BRET, etc. Broadly, the results presented here highlight the role of collective dynamics in enzyme activity and that mutations in an enzyme can be selected based on their contribution in the collective dynamics of the protein.

## Methods

### Structural modeling of WT and E50A mutant picALuc

We utilized the available nuclear magnetic resonance (NMR)-derived structure of GLuc (PDB: 7D2O)^1^ to generate a homology-based structural model of the picALuc using the SWISS-MODEL webserver^51^.^41, 44, 52^ The two proteins, picALuc and GLuc show considerable sequence similarity (92.5%; and 85% identity) with key differences in the N- and C-terminal (Supplementary Figure 1A) allowing high confidence structural modelling of picALuc. Notably, picALuc lacks N-terminal a-helices 1 and 2 and some residues in the C-terminal region of ALuc (Supplementary Figure 1B). Quality of the modelled structure was assessed using the MolProbity software (version 4.4) ^53^ available as a part of the SWISS-MODEL webserver with a score of 2.90 and with only two Gly residues (located in the long loop between α-helices 3 and 4) out of a total of 120 residues (1.69%; the GLuc template structure 7D2O^1^ showed 4 out 168 residues, which is equal to 2.8%) as Ramachandran plot outliers. Importantly, all disulfide bridges observed in GLuc structure were maintained in the modelled picALuc structure (Supplementary Figure 1C). Structural model of the E50A mutant picALuc was generated by backbone rotamer-dependent replacement of the residue E with an A using Pymol (The PyMOL Molecular Graphics System, Version 2.0.0, Schrödinger, LLC; pymol.org; New York, NY, USA).

### MD Simulations

MD simulations of the modelled WT and E50A mutant picALuc structures was performed essentially as described previously.^31, 44^ Inputs files including topology and parameter files were prepared using the QwikMD plugin^54^ available in the Visual Molecular Dynamics (VMD) 1.9.3 software^55^ and simulations were performed using the Nanoscale Molecular Dynamics (NAMD) 2.13 software^35^ and CHARMM36 force field^56^. Briefly, proteins were solvated in explicit solvent using TIP3P^57^ cubic water box containing 0.15 M NaCl for charge neutralization and periodic boundary conditions applied with a 2 fs integration time-step. Prior to production simulations, energy minimization was performed on each system for 2000 steps (4 ps).^44^ Subsequently, the systems were graduated heated from 60 K to 310 K at 1 K interval to reach the 310 K equilibrium temperature. Following thermalization, temperature was maintained at 310 K using Langevin temperature control and pressure was maintained at 1.0 atm using Nose-Hoover Langevin piston pressure control^58^. The systems were then equilibrated for 500,000 steps (1 ns). GaMD simulations were then run using the integrated GaMD module in NAMD and its default parameters.^36, 37^ This included a 2 ns of conventional molecular dynamics equilibration run for collecting potential statistics that were used for calculating acceleration parameters, and another 50 ns equilibration run with the addition of boost potential^37, 59^, and finally GaMD production runs for 1,000 ns. A 400 ps preparatory runs were included before each of the equilibration steps in GaMD. All GaMD simulations were run at the “dual-boost” level with one boost potential applied to the dihedral energy term and the other applied to the total potential energy term^36, 60, 61^ with a standard deviation upper limit set to 6.0 kcal/mol. Short-range non-bonded interactions were defined at 12 Å cut-off with 10 Å switching distance, while Particle-mesh Ewald (PME) scheme was used to handle long-range electrostatic interactions at 1 Å PME grid spacing. Simulation frames were saved every 10,000 steps (20 ps). Simulation trajectories were analyzed using available tools in the Visual Molecular Dynamics (VMD) software^55^ and the MDAnalysis package^46, 47^. Backbone root-mean-square deviation (RMSD), root-mean-square fluctuation (RMSF), solvent accessible surface area (SASA), radius of gyration (RoG), energy calculations, secondary structure salt bridges and H-bond occupancy were performed using VMD^55, 62, 63^. Pairwise RMSD, number of H-bonds formed and S atom distances in disulfide bridges were calculated using the MDAnalysis package^46, 47^. Dynamic cross-correlation (DCC) analysis of the trajectories were performed using the calc_correlation.py script available as a part of the MD-TASK suite of MD trajectory analysis software^43^ over the whole 1 μs trajectory with a step size of 2 (alternating frames). Principal component analysis (PCA) was performed using the PCA module available in the MDAnalysis package^46, 47^. Contribution of individual residues to PC1 and 2 were calculated by projecting the principal components on Cα atoms of the proteins using modules in the MDAnalysis package^46, 47^ followed by calculation of individual Cα atom displacement (root mean squared). Movies were prepared from 500 frames out of the 50,000 frames of the 1 μs trajectory (with a step size of 100 frames) generated using the VMD Movie Maker plugin^55^ and compiled at 20 fps using the Fiji distribution of ImageJ software ^64^. All structural images were generated using Pymol (The PyMOL Molecular Graphics System, Version 2.0.0, Schrödinger, LLC; pymol.org; New York, NY, USA).

### mGL-picALuc plasmid construct generation

Complete nucleotide and amino acid sequences of all constructs designed and utilized in the current manuscript have been included in the Supplementary Text. A codon optimized gene sequence of picALuc generated using the previously described amino acid sequence^30^ was synthesized and subcloned into the pcDNA3.1-mGL-NLuc plasmid (generated by synthesizing the mGL gene and subcloned into the previously described pmNG-Mpro-Nter-auto-NLuc plasmid using *Eco*RI-*Bam*HI sites) using *Bam*HI and *Xho*I restriction enzyme sites (GenScript, Singapore) to generate the pcDNA3.1-mGL-picALuc plasmid. Additionally, the SARS-CoV-2 M^pro^ N-terminal autocleavage sequence was included in the N-terminal^31, 65^ and 3x FLAG-tag sequence was included at the C-terminal. Glu10Ala (E10A), Glu50Ala (E50A) and Asp94Ala (D94A) mutations were generated in the pcDNA3.1-mGL-picALuc plasmid (GenScript, Singapore). For generating the picSm and picSm-GCN4 fragment expressing plasmids, nucleotide sequences encoding picALuc N-terminal helixα1 (K1 to A22) or one with the GCN4 peptide (EELLSKNYHLENEVARLKK) were synthesized and subcloned into the pcDNA3.1-mGL-NLuc plasmid using *Bam*HI-*Xho*I sites (GenScript, Singapore). For generating the picLg and picLg-GCN4 fragment expressing plasmids, nucleotide sequences encoding picALuc residues G23-C120 or with the GCN4 peptide were synthesized and subcloned into pcDNA3.1-mGL-NLuc plasmid using the *Hind*III-*Xho*I sites (GenScript, Singapore).

### Cell culture and transfection

HEK 293T cells were cultured in Dulbecco’s Modified Eagle Medium (DMEM) supplemented with 10% fetal bovine serum, and 1% penicillin-streptomycin and grown at 37 °C in 5% CO_2_. ^34, 66, 67, 68, 69, 70, 71^ Transfections were performed using polyethyleneimine (PEI) lipid according to the manufacturers’ protocol. Briefly, HEK 293T cells were seeded onto 96-well white plates before 24 h of transfection. The plasmid DNA (encoding either the WT or mutant picALucs; 125 ng/well), Opti-MEM (Invitrogen; 31985088; 25 μL/well) and PEI lipid (Sigma-Aldrich; 408727-100mL; 0.625 μL/well) were mixed and incubated at room temperature for 30 min prior to addition to the cells. For in vitro biochemical and thermal stability assays, cells were transfected in 60 mm dishes using appropriate proportion of DNA, PEI lipid and Opti-MEM. For the protein fragment complementation assays, HEK 293T cells were transfected with a range of picSm or picSm-GCN4 and picLg or picLg-GCN4 plasmid DNA ratios with picLg or picLg-GCN4 maintained at 25 ng/well while total plasmid concentrations were maintained at 150 ng/well using a control plasmid (pcDNA3-HA). The PEI stock solution of 2 mg/mL was prepared by diluting in sterile Milli-Q water and stored at −80 °C for subsequent usage.

### Live cell assays

Live cell assays to determine fluorescence, bioluminescence and bioluminescence emission spectra were performed by transfecting HEK 293T cells with either the WT or the mutant pcDNA3.1-mGL-picALuc plasmid constructs.^68, 69, 70, 71^ For SARS-CoV-2 M^pro^-mediated cleavage of mGL-picALuc, cells transfected with pcDNA3.1-mGL-picALuc plasmid along with either pLVX-EF1alpha-SARS-CoV-2-nsp5-2xStrep-IRES-Puro (M^pro^ WT) (a gift from Nevan Krogan (Addgene plasmid # 141370; http://n2t.net/addgene:141370 ; RRID:Addgene_141370)^72^ or pLVX-EF1alpha-SARS-CoV-2-nsp5-C145A-2xStrep-IRES-Puro (C145A mutant M^pro^) plasmid (a gift from Nevan Krogan (Addgene plasmid # 141371; http://n2t.net/addgene:141371; RRID:Addgene_141371)^72^. For the picALuc-based protein complementation assay, cells were transfected with the indicated ratios of plasmid DNA amounts. After 48 h of transfection, mGL fluorescence were measured using a Tecan SPARK® multimode microplate reader prior to bioluminescence measurements. Samples were excited at a wavelength 480 nm and emission acquired at a wavelength of 530 nm. Bioluminescence was measured in the cells after addition of luciferase substrate, coelenterazine h, addition at a final concentration of 5 μM. Bioluminescence spectra was acquired using the Tecan SPARK® multimode microplate reader between 380 to 665 nm wavelengths with an acquisition time of 400 ms for each wavelength. Bioluminescence spectral data was normalized by dividing emissions at all wavelengths with that of emission at 483 nm. BRET was calculated as a ratio of emission at 516 nm (corresponding to the emission maxima of acceptor mGL) and 483 nm (corresponding to the emission maxima of donor picALuc).^31, 32, 33, 40, 41, 42^ Bioluminescence of each sample was calculated from emissions at wavelengths between 380 and 665 nm and was first normalized with mGL fluorescence (relative to WT values) and then with that of bioluminescence of WT picALuc. For protein complementation assay, total bioluminescence (all wavelengths) was measured in all samples following mGL fluorescence measurements. All experiments were performed thrice in triplicates.

### In vitro assays

In vitro assays were performed using lysates prepared from cells expressing the respective picALuc constructs as described previously.^32, 33, 40, 41, 42^ After 48 h of transfection, cells were washed thrice in chilled Dulbecco’s phosphate-buffered saline (DPBS) and harvested and lysed by sonication on ice in a buffer containing 50 mM HEPES (pH 7.5), 100 mM NaCl, 2 mM EDTA, 1 mM dithiothreitol (DTT), 1× protease inhibitor cocktail (ThermoFisher Scientific, Massachusetts, USA) and 10% glycerol. Sonicated samples were centrifuged at 4 ͦC for 1 h and supernatant was collected for further experiments.

Biochemical experiments to determine substrate affinity and reaction velocity, equivalent amounts of the proteins were taken after normalization with mGL fluorescence and incubated with a range of coelenterazine h concentrations. Data were fit to a allosteric sigmoidal model considering bioluminescence output (counts per second) representing the rate of reaction to determine *V*_max_ and *K*_half_ values. Thermal stability of the proteins was determined by measuring bioluminescence from equivalent amounts of cell lysates after incubation at a range of temperatures for 10 min. Data were fit to a Boltzmann sigmoidal model to determine the temperature at which the protein shows half maximal activity (*T*_m_). Bioluminescence was measured after addition of the substrate in a GloMax® Discover Microplate Reader (Promega). All experiments were performed thrice in triplicates.

### Data Analysis and Figure Preparation

MATLAB (release 2021b), Matplotlib, GraphPad Prism (version 9 for macOS, GraphPad Software, La Jolla California USA; www.graphpad.com) and Microsoft Excel were used for data analysis and graph preparation. Figures were assembled using Adobe Illustrator.

## Acknowledgements

This work is supported by an internal funding from the College of Health & Life Sciences, Hamad Bin Khalifa University, a member of the Qatar Foundation. Some of the computational research work reported in the manuscript were performed using high-performance computer resources and services provided by the Research Computing group in Texas A&M University at Qatar. Research Computing is funded by the Qatar Foundation for Education, Science and Community Development (http://www.qf.org.qa).

## Author Contributions

K.H.B. conceived and performed the experiments, analyzed data, prepared figure panels, wrote, and approved the manuscript.

