## Supplementary for "A Brighter picALuc Generated Through the Loss of a Salt Bridge Interaction"

Kabir H Biswas[1,*]

**Affiliation**:
[1]Division of Biological and Biomedical Sciences, College of Health & Life Sciences, Hamad Bin Khalifa University, Qatar Foundation, Doha – 34110, Qatar

ORCID:
Kabir H Biswas: 0000-0001-9194-4127

### Supplementary Text

##### *mGL-picALuc nucleotide sequence:*

ATGGGAAGTTCACATCATCATCATCATCACTCATCAGGACTGGTGCCACGGGGGTCTGAATTCGGCATGGTGAGCAAGGGCGAGGAGCTGTTCACCGGGGTGGTGCCCATCC
TGGTCGAGCTGGACGGCGACGTAAACGGCCACAAGTTCAGCGTCCGCGGCGAGGGCGAGGGCGATGCCACCAACGGCAAGCTGACCCTGAAGTTCATCTGCACCACCGGCAA
GCTGCCCGTGCCCTGGCCCACCCTCGTGACCACCTTAGGCTACGGCGTGGCCTGCTTCGCCCGCTACCCCGACCACATGAAGCAGCACGACTTCTTCAAGTCCGCCATGCCC
GAAGGCTACGTCCAGGAGCGCACCATCTCTTTCAAGGACGACGGTACCTACAAGACCCGCGCCGAGGTGAAGTTCGAGGGCGACACCCTGGTGAACCGCATCGTGCTGAAGG
GCATCGACTTCAAGGAGGACGGCAACATCCTGGGGCACAAGCTGGAGTACAACTTCAACAGCCACAAGGTCTATATCACGGCCGACAAGCAGAAGAACGGCATCAAGGCTAA
CTTCAAGACCCGCCACAACGTTGAGGACGGCGGCGTGCAGCTCGCCGACCACTACCAGCAGAACACCCCCATCGGCGACGGCCCCGTGCTGCTGCCCGACAACCACTACCTG
AGCCATCAGTCCAAACTGAGCAAAGACCCCAACGAGAAGCGCGATCACATGGTCCTGAAGGAGAGGGTGACCGCCGCCGGGATTACACATGACATGGACGAGCTGTACAAGT
ACGGATCCGCGGCCGCCACCGAGAACCTGTATGCAGTGCTCCAAAGCGGATTTCGCGGCTCTGGCAGCGCTATGAAGCTGCCCGGCAAGAAGCTGCCCCTGGAGGTGCTGAA
GGAGCTGGAGGCCAACGCCCAGAAGGCCGGCTGCACCAGGGGCTGCCTGATCTGCCTGAGCCACATCAAGTGCACCGCCAAGATGAAGAAGTGGCTGCCCGGCAGGTGCGAG
AGCTGGGAGGGCGACAAGGAGACCGGCCAGGGCGGCATCGGCGAGGCCATCGTGGACATCCCCGAGATCCCCGGCTTCAAGGAGCTGGCCCCCATGGAGCAGTTCATCGCCC
AGGTGGACCTGTGCGCCGACTGCACCACCGGCTGCCTGAAGGGCCTGGCCAACGTGAAGTGCAGCGCCCTGCTGAAGAAGTGGCTGCCCAGCAGGTGCGGTACCGACTACAA
AGACCATGACGGTGATTATAAAGATCATGACATCGATTACAAGGATGACGATGACAAGGATATCTGA

##### *mGL-picALuc amino acid sequence:*

MGSSHHHHHHSSGLVPRGSEFGMVSKGEELFTGVVPILVELDGDVNGHKFSVRGEGEGDATNGKLTLKFICTTGKLPVPWPTLVTTLGYGVACFARYPDHMKQHDFFKSAMP
EGYVQERTISFKDDGTYKTRAEVKFEGDTLVNRIVLKGIDFKEDGNILGHKLEYNFNSHKVYITADKQKNGIKANFKTRHNVEDGGVQLADHYQQNTPIGDGPVLLPDNHYL
SHQSKLSKDPNEKRDHMVLKERVTAAGITHDMDELYKYGSAAATENLYAVLQSGFRGSGSAMKLPGKKLPLEVLKELEANAQKAGCTRGCLICLSHIKCTAKMKKWLPGRCE
SWEGDKETGQGGIGEAIVDIPEIPGFKELAPMEQFIAQVDLCADCTTGCLKGLANVKCSALLKKWLPSRCGTDYKDHDGDYKDHDIDYKDDDDKDI*

##### *mGL-picALuc(E10A) nucleotide sequence:*

ATGGGAAGTTCACATCATCATCATCATCACTCATCAGGACTGGTGCCACGGGGGTCTGAATTCGGCATGGTGAGCAAGGGCGAGGAGCTGTTCACCGGGGTGGTGCCCATCC
TGGTCGAGCTGGACGGCGACGTAAACGGCCACAAGTTCAGCGTCCGCGGCGAGGGCGAGGGCGATGCCACCAACGGCAAGCTGACCCTGAAGTTCATCTGCACCACCGGCAA
GCTGCCCGTGCCCTGGCCCACCCTCGTGACCACCTTAGGCTACGGCGTGGCCTGCTTCGCCCGCTACCCCGACCACATGAAGCAGCACGACTTCTTCAAGTCCGCCATGCCC
GAAGGCTACGTCCAGGAGCGCACCATCTCTTTCAAGGACGACGGTACCTACAAGACCCGCGCCGAGGTGAAGTTCGAGGGCGACACCCTGGTGAACCGCATCGTGCTGAAGG
GCATCGACTTCAAGGAGGACGGCAACATCCTGGGGCACAAGCTGGAGTACAACTTCAACAGCCACAAGGTCTATATCACGGCCGACAAGCAGAAGAACGGCATCAAGGCTAA
CTTCAAGACCCGCCACAACGTTGAGGACGGCGGCGTGCAGCTCGCCGACCACTACCAGCAGAACACCCCCATCGGCGACGGCCCCGTGCTGCTGCCCGACAACCACTACCTG
AGCCATCAGTCCAAACTGAGCAAAGACCCCAACGAGAAGCGCGATCACATGGTCCTGAAGGAGAGGGTGACCGCCGCCGGGATTACACATGACATGGACGAGCTGTACAAGT
ACGGATCCGCGGCCGCCACCGAGAACCTGTATGCAGTGCTCCAAAGCGGATTTCGCGGCTCTGGCAGCGCTATGAAGCTGCCCGGCAAGAAGCTGCCCCTGGCCGTGCTGAA
GGAGCTGGAGGCCAACGCCCAGAAGGCCGGCTGCACCAGGGGCTGCCTGATCTGCCTGAGCCACATCAAGTGCACCGCCAAGATGAAGAAGTGGCTGCCCGGCAGGTGCGAG
AGCTGGGAGGGCGACAAGGAGACCGGCCAGGGCGGCATCGGCGAGGCCATCGTGGACATCCCCGAGATCCCCGGCTTCAAGGAGCTGGCCCCCATGGAGCAGTTCATCGCCC
AGGTGGACCTGTGCGCCGACTGCACCACCGGCTGCCTGAAGGGCCTGGCCAACGTGAAGTGCAGCGCCCTGCTGAAGAAGTGGCTGCCCAGCAGGTGCGGTACCGACTACAA
AGACCATGACGGTGATTATAAAGATCATGACATCGATTACAAGGATGACGATGACAAGGATATCTGA

##### *mGL-picALuc(E10A) amino acid sequence:*

MGSSHHHHHHSSGLVPRGSEFGMVSKGEELFTGVVPILVELDGDVNGHKFSVRGEGEGDATNGKLTLKFICTTGKLPVPWPTLVTTLGYGVACFARYPDHMKQHDFFKSAMP
EGYVQERTISFKDDGTYKTRAEVKFEGDTLVNRIVLKGIDFKEDGNILGHKLEYNFNSHKVYITADKQKNGIKANFKTRHNVEDGGVQLADHYQQNTPIGDGPVLLPDNHYL
SHQSKLSKDPNEKRDHMVLKERVTAAGITHDMDELYKYGSAAATENLYAVLQSGFRGSGSAMKLPGKKLPLAVLKELEANAQKAGCTRGCLICLSHIKCTAKMKKWLPGRCE
SWEGDKETGQGGIGEAIVDIPEIPGFKELAPMEQFIAQVDLCADCTTGCLKGLANVKCSALLKKWLPSRCGTDYKDHDGDYKDHDIDYKDDDDKDI*

##### *mGL-picALuc(E50A) nucleotide sequence:*

ATGGGAAGTTCACATCATCATCATCATCACTCATCAGGACTGGTGCCACGGGGGTCTGAATTCGGCATGGTGAGCAAGGGCGAGGAGCTGTTCACCGGGGTGGTGCCCATCC
TGGTCGAGCTGGACGGCGACGTAAACGGCCACAAGTTCAGCGTCCGCGGCGAGGGCGAGGGCGATGCCACCAACGGCAAGCTGACCCTGAAGTTCATCTGCACCACCGGCAA
GCTGCCCGTGCCCTGGCCCACCCTCGTGACCACCTTAGGCTACGGCGTGGCCTGCTTCGCCCGCTACCCCGACCACATGAAGCAGCACGACTTCTTCAAGTCCGCCATGCCC
GAAGGCTACGTCCAGGAGCGCACCATCTCTTTCAAGGACGACGGTACCTACAAGACCCGCGCCGAGGTGAAGTTCGAGGGCGACACCCTGGTGAACCGCATCGTGCTGAAGG
GCATCGACTTCAAGGAGGACGGCAACATCCTGGGGCACAAGCTGGAGTACAACTTCAACAGCCACAAGGTCTATATCACGGCCGACAAGCAGAAGAACGGCATCAAGGCTAA
CTTCAAGACCCGCCACAACGTTGAGGACGGCGGCGTGCAGCTCGCCGACCACTACCAGCAGAACACCCCCATCGGCGACGGCCCCGTGCTGCTGCCCGACAACCACTACCTG
AGCCATCAGTCCAAACTGAGCAAAGACCCCAACGAGAAGCGCGATCACATGGTCCTGAAGGAGAGGGTGACCGCCGCCGGGATTACACATGACATGGACGAGCTGTACAAGT
ACGGATCCGCGGCCGCCACCGAGAACCTGTATGCAGTGCTCCAAAGCGGATTTCGCGGCTCTGGCAGCGCTATGAAGCTGCCCGGCAAGAAGCTGCCCCTGGAGGTGCTGAA
GGAGCTGGAGGCCAACGCCCAGAAGGCCGGCTGCACCAGGGGCTGCCTGATCTGCCTGAGCCACATCAAGTGCACCGCCAAGATGAAGAAGTGGCTGCCCGGCAGGTGCGCC
AGCTGGGAGGGCGACAAGGAGACCGGCCAGGGCGGCATCGGCGAGGCCATCGTGGACATCCCCGAGATCCCCGGCTTCAAGGAGCTGGCCCCCATGGAGCAGTTCATCGCCC
AGGTGGACCTGTGCGCCGACTGCACCACCGGCTGCCTGAAGGGCCTGGCCAACGTGAAGTGCAGCGCCCTGCTGAAGAAGTGGCTGCCCAGCAGGTGCGGTACCGACTACAA
AGACCATGACGGTGATTATAAAGATCATGACATCGATTACAAGGATGACGATGACAAGGATATCTGA

##### *mGL-picALuc(E50A) amino acid sequence:*

MGSSHHHHHHSSGLVPRGSEFGMVSKGEELFTGVVPILVELDGDVNGHKFSVRGEGEGDATNGKLTLKFICTTGKLPVPWPTLVTTLGYGVACFARYPDHMKQHDFFKSAMP
EGYVQERTISFKDDGTYKTRAEVKFEGDTLVNRIVLKGIDFKEDGNILGHKLEYNFNSHKVYITADKQKNGIKANFKTRHNVEDGGVQLADHYQQNTPIGDGPVLLPDNHYL
SHQSKLSKDPNEKRDHMVLKERVTAAGITHDMDELYKYGSAAATENLYAVLQSGFRGSGSAMKLPGKKLPLEVLKELEANAQKAGCTRGCLICLSHIKCTAKMKKWLPGRCA
SWEGDKETGQGGIGEAIVDIPEIPGFKELAPMEQFIAQVDLCADCTTGCLKGLANVKCSALLKKWLPSRCGTDYKDHDGDYKDHDIDYKDDDDKDI*

##### *mGL-picALuc(D94A) nucleotide sequence:*

ATGGGAAGTTCACATCATCATCATCATCACTCATCAGGACTGGTGCCACGGGGGTCTGAATTCGGCATGGTGAGCAAGGGCGAGGAGCTGTTCACCGGGGTGGTGCCCATCC
TGGTCGAGCTGGACGGCGACGTAAACGGCCACAAGTTCAGCGTCCGCGGCGAGGGCGAGGGCGATGCCACCAACGGCAAGCTGACCCTGAAGTTCATCTGCACCACCGGCAA
GCTGCCCGTGCCCTGGCCCACCCTCGTGACCACCTTAGGCTACGGCGTGGCCTGCTTCGCCCGCTACCCCGACCACATGAAGCAGCACGACTTCTTCAAGTCCGCCATGCCC
GAAGGCTACGTCCAGGAGCGCACCATCTCTTTCAAGGACGACGGTACCTACAAGACCCGCGCCGAGGTGAAGTTCGAGGGCGACACCCTGGTGAACCGCATCGTGCTGAAGG
GCATCGACTTCAAGGAGGACGGCAACATCCTGGGGCACAAGCTGGAGTACAACTTCAACAGCCACAAGGTCTATATCACGGCCGACAAGCAGAAGAACGGCATCAAGGCTAA
CTTCAAGACCCGCCACAACGTTGAGGACGGCGGCGTGCAGCTCGCCGACCACTACCAGCAGAACACCCCCATCGGCGACGGCCCCGTGCTGCTGCCCGACAACCACTACCTG
AGCCATCAGTCCAAACTGAGCAAAGACCCCAACGAGAAGCGCGATCACATGGTCCTGAAGGAGAGGGTGACCGCCGCCGGGATTACACATGACATGGACGAGCTGTACAAGT
ACGGATCCGCGGCCGCCACCGAGAACCTGTATGCAGTGCTCCAAAGCGGATTTCGCGGCTCTGGCAGCGCTATGAAGCTGCCCGGCAAGAAGCTGCCCCTGGAGGTGCTGAA
GGAGCTGGAGGCCAACGCCCAGAAGGCCGGCTGCACCAGGGGCTGCCTGATCTGCCTGAGCCACATCAAGTGCACCGCCAAGATGAAGAAGTGGCTGCCCGGCAGGTGCGAG
AGCTGGGAGGGCGACAAGGAGACCGGCCAGGGCGGCATCGGCGAGGCCATCGTGGACATCCCCGAGATCCCCGGCTTCAAGGAGCTGGCCCCCATGGAGCAGTTCATCGCCC
AGGTGGACCTGTGCGCCGCCTGCACCACCGGCTGCCTGAAGGGCCTGGCCAACGTGAAGTGCAGCGCCCTGCTGAAGAAGTGGCTGCCCAGCAGGTGCGGTACCGACTACAA
AGACCATGACGGTGATTATAAAGATCATGACATCGATTACAAGGATGACGATGACAAGGATATCTGA

***mGL-picALuc(D94A) amino acid sequence:***

MGSSHHHHHHSSGLVPRGSEFGMVSKGEELFTGVVPILVELDGDVNGHKFSVRGEGEGDATNGKLTLKFICTTGKLPVPWPTLVTTLGYGVACFARYPDHMKQHDFFKSAMP
EGYVQERTISFKDDGTYKTRAEVKFEGDTLVNRIVLKGIDFKEDGNILGHKLEYNFNSHKVYITADKQKNGIKANFKTRHNVEDGGVQLADHYQQNTPIGDGPVLLPDNHYL
SHQSKLSKDPNEKRDHMVLKERVTAAGITHDMDELYKYGSAAATENLYAVLQSGFRGSGSAMKLPGKKLPLEVLKELEANAQKAGCTRGCLICLSHIKCTAKMKKWLPGRCE
SWEGDKETGQGGIGEAIVDIPEIPGFKELAPMEQFIAQVDLCAACTTGCLKGLANVKCSALLKKWLPSRCGTDYKDHDGDYKDHDIDYKDDDDKDI*

***mGL-picSm nucleotide sequence:***

ATGGGAAGTTCACATCATCATCATCATCACTCATCAGGACTGGTGCCACGGGGGTCTGAATTCGGCATGGTGAGCAAGGGCGAGGAGCTGTTCACCGGGGTGGTGCCCATCC
TGGTCGAGCTGGACGGCGACGTAAACGGCCACAAGTTCAGCGTCCGCGGCGAGGGCGAGGGCGATGCCACCAACGGCAAGCTGACCCTGAAGTTCATCTGCACCACCGGCAA
GCTGCCCGTGCCCTGGCCCACCCTCGTGACCACCTTAGGCTACGGCGTGGCCTGCTTCGCCCGCTACCCCGACCACATGAAGCAGCACGACTTCTTCAAGTCCGCCATGCCC
GAAGGCTACGTCCAGGAGCGCACCATCTCTTTCAAGGACGACGGTACCTACAAGACCCGCGCCGAGGTGAAGTTCGAGGGCGACACCCTGGTGAACCGCATCGTGCTGAAGG
GCATCGACTTCAAGGAGGACGGCAACATCCTGGGGCACAAGCTGGAGTACAACTTCAACAGCCACAAGGTCTATATCACGGCCGACAAGCAGAAGAACGGCATCAAGGCTAA
CTTCAAGACCCGCCACAACGTTGAGGACGGCGGCGTGCAGCTCGCCGACCACTACCAGCAGAACACCCCCATCGGCGACGGCCCCGTGCTGCTGCCCGACAACCACTACCTG
AGCCATCAGTCCAAACTGAGCAAAGACCCCAACGAGAAGCGCGATCACATGGTCCTGAAGGAGAGGGTGACCGCCGCCGGGATTACACATGACATGGACGAGCTGTACAAGT
ACGGATCCGCGGCCGCCACCGAGAACCTGTATGCAGTGCTCCAAAGCGGATTTCGCGGCTCTGGCAGCGCTATGAAGCTGCCCGGCAAGAAGCTGCCCCTGGAGGTGCTGAA
GGAGCTGGAGGCCAACGCCCAGAAGGCCTGA

***mGL-picSm amino acid sequence:***

MGSSHHHHHHSSGLVPRGSEFGMVSKGEELFTGVVPILVELDGDVNGHKFSVRGEGEGDATNGKLTLKFICTTGKLPVPWPTLVTTLGYGVACFARYPDHMKQHDFFKSAMP
EGYVQERTISFKDDGTYKTRAEVKFEGDTLVNRIVLKGIDFKEDGNILGHKLEYNFNSHKVYITADKQKNGIKANFKTRHNVEDGGVQLADHYQQNTPIGDGPVLLPDNHYL
SHQSKLSKDPNEKRDHMVLKERVTAAGITHDMDELYKYGSAAATENLYAVLQSGFRGSGSAMKLPGKKLPLEVLKELEANAQKA*

***mGL-picSm-GNC4 nucleotide sequence:***

ATGGGAAGTTCACATCATCATCATCATCACTCATCAGGACTGGTGCCACGGGGGTCTGAATTCGGCATGGTGAGCAAGGGCGAGGAGCTGTTCACCGGGGTGGTGCCCATCC
TGGTCGAGCTGGACGGCGACGTAAACGGCCACAAGTTCAGCGTCCGCGGCGAGGGCGAGGGCGATGCCACCAACGGCAAGCTGACCCTGAAGTTCATCTGCACCACCGGCAA
GCTGCCCGTGCCCTGGCCCACCCTCGTGACCACCTTAGGCTACGGCGTGGCCTGCTTCGCCCGCTACCCCGACCACATGAAGCAGCACGACTTCTTCAAGTCCGCCATGCCC
GAAGGCTACGTCCAGGAGCGCACCATCTCTTTCAAGGACGACGGTACCTACAAGACCCGCGCCGAGGTGAAGTTCGAGGGCGACACCCTGGTGAACCGCATCGTGCTGAAGG
GCATCGACTTCAAGGAGGACGGCAACATCCTGGGGCACAAGCTGGAGTACAACTTCAACAGCCACAAGGTCTATATCACGGCCGACAAGCAGAAGAACGGCATCAAGGCTAA
CTTCAAGACCCGCCACAACGTTGAGGACGGCGGCGTGCAGCTCGCCGACCACTACCAGCAGAACACCCCCATCGGCGACGGCCCCGTGCTGCTGCCCGACAACCACTACCTG
AGCCATCAGTCCAAACTGAGCAAAGACCCCAACGAGAAGCGCGATCACATGGTCCTGAAGGAGAGGGTGACCGCCGCCGGGATTACACATGACATGGACGAGCTGTACAAGT
ACGGATCCGCGGCCGCCACCGAGAACCTGTATGCAGTGCTCCAAAGCGGATTTCGCGGCTCTGGCAGCGCTATGAAGCTGCCCGGCAAGAAGCTGCCCCTGGAGGTGCTGAA
GGAGCTGGAGGCCAACGCCCAGAAGGCCGGCTCTGGCTCTATCGATGGCTCTGGCTCTGAAGAACTGCTGAGCAAAAACTATCATCTGGAAAACGAAGTGGCGCGCCTGAAA
AAACTGGTGGGCGAACGCTGA

***mGL-picSm-GCN4 amino acid sequence:***

MGSSHHHHHHSSGLVPRGSEFGMVSKGEELFTGVVPILVELDGDVNGHKFSVRGEGEGDATNGKLTLKFICTTGKLPVPWPTLVTTLGYGVACFARYPDHMKQHDFFKSAMP
EGYVQERTISFKDDGTYKTRAEVKFEGDTLVNRIVLKGIDFKEDGNILGHKLEYNFNSHKVYITADKQKNGIKANFKTRHNVEDGGVQLADHYQQNTPIGDGPVLLPDNHYL
SHQSKLSKDPNEKRDHMVLKERVTAAGITHDMDELYKYGSAAATENLYAVLQSGFRGSGSAMKLPGKKLPLEVLKELEANAQKAGSGSIDGSGSEELLSKNYHLENEVARLK
KLVGER*

***picLg nucleotide sequence:***

ATGGGCTGCACCAGGGGCTGCCTGATCTGCCTGAGCCACATCAAGTGCACCGCCAAGATGAAGAAGTGGCTGCCCGGCAGGTGCGAGAGCTGGGAGGGCGACAAGGAGACCG
GCCAGGGCGGCATCGGCGAGGCCATCGTGGACATCCCCGAGATCCCCGGCTTCAAGGAGCTGGCCCCCATGGAGCAGTTCATCGCCCAGGTGGACCTGTGCGCCGACTGCAC
CACCGGCTGCCTGAAGGGCCTGGCCAACGTGAAGTGCAGCGCCCTGCTGAAGAAGTGGCTGCCCAGCAGGTGCGGTACCGACTACAAAGACCATGACGGTGATTATAAAGAT
CATGACATCGATTACAAGGATGACGATGACAAGGGATCCTTAAGGATATCTGAGCGGCCGCGAATTCCTCGAGTCTAG

***picLg amino acid sequence:***

MGCTRGCLICLSHIKCTAKMKKWLPGRCESWEGDKETGQGGIGEAIVDIPEIPGFKELAPMEQFIAQVDLCADCTTGCLKGLANVKCSALLKKWLPSRCGTDYKDHDGDYKD
HDIDYKDDDDKGSLRISERPRIPRV*

***picLg-GNC4 nucleotide sequence:***

ATGGGCTGCACCAGGGGCTGCCTGATCTGCCTGAGCCACATCAAGTGCACCGCCAAGATGAAGAAGTGGCTGCCCGGCAGGTGCGAGAGCTGGGAGGGCGACAAGGAGACCG
GCCAGGGCGGCATCGGCGAGGCCATCGTGGACATCCCCGAGATCCCCGGCTTCAAGGAGCTGGCCCCCATGGAGCAGTTCATCGCCCAGGTGGACCTGTGCGCCGACTGCAC
CACCGGCTGCCTGAAGGGCCTGGCCAACGTGAAGTGCAGCGCCCTGCTGAAGAAGTGGCTGCCCAGCAGGTGCGGTACCGGCTCTGGCTCTGGCTCTGGCTCTGGCTCTGAA
GAACTGCTGAGCAAAAACTATCATCTGGAAAACGAAGTGGCGCGCCTGAAAAAACTGGTGGGCGAACGCGACTACAAAGACCATGACGGTGATTATAAAGATCATGACATCG
ATTACAAGGATGACGATGACAAGGGATCCTTAAGGATATCTGAGCGGCCGCGAATTCCTCGAGTCTAG

***picLg-GCN4 amino acid sequence:***

MGCTRGCLICLSHIKCTAKMKKWLPGRCESWEGDKETGQGGIGEAIVDIPEIPGFKELAPMEQFIAQVDLCADCTTGCLKGLANVKCSALLKKWLPSRCGTGSGSGSGSGSE
ELLSKNYHLENEVARLKKLVGERDYKDHDGDYKDHDIDYKDDDDKGSLRISERPRIPRV*

*Color code:*
Green: mGreenLantern (mGL)
Grey: SARS-CoV-2 Mpro cleavage sequence
Blue: picALuc
Yellow: 3xFLAG-tag
Magent: GCN4 peptide
Mutated residues are highlighted in light red color

#### Supplementary Table

***Supplementary Table 1. H-bond occupancy table.*** *H-bonds showing >5% occupancy. H-bonds formed Glu50 side chain are highlighted in green while that formed by its main chain is highlighted in light green. Similarly, H-bonds formed by Glu10 side chain are highlighted in blue while that formed by its main chain is highlighted in light blue. H-bonds formed by Asp94 side chain is highlighted in grey.*

| # | Donor | Acceptor | Occupancy |
|---|---|---|---|
| 1 | LYS36-Side | ASP55-Side | 91.31% |
| 2 | ARG119-Side | GLU53-Side | 75.32% |
| 3 | LYS42-Side | GLU50-Side | 68.80% |
| 4 | GLU16-Main | LEU12-Main | 65.42% |
| 5 | ASN18-Main | GLU14-Main | 62.57% |
| 6 | LEU12-Main | PRO8-Main | 60.13% |
| 7 | LYS13-Main | LEU9-Main | 57.13% |
| 8 | LEU15-Main | VAL11-Main | 55.16% |
| 9 | ARG26-Side | GLU16-Side | 52.80% |
| 10 | LYS101-Main | ALA22-Main | 51.20% |
| 11 | ALA19-Main | LEU15-Main | 48.66% |
| 12 | SER109-Side | GLU14-Side | 44.72% |
| 13 | LEU45-Main | MET41-Main | 44.70% |
| 14 | LEU32-Main | CYS28-Main | 44.50% |
| 15 | SER33-Side | LEU29-Main | 41.72% |
| 16 | CYS24-Main | ALA19-Main | 40.07% |
| 17 | TRP52-Side | LYS5-Main | 38.88% |
| 18 | ALA22-Main | ASN18-Main | 37.40% |
| 19 | GLN20-Main | GLU16-Main | 36.60% |
| 20 | LYS42-Main | THR38-Main | 35.86% |
| 21 | LYS113-Side | CYS120-Side | 33.52% |
| 22 | ALA17-Main | LYS13-Main | 33.06% |
| 23 | LYS21-Main | ALA17-Main | 33.02% |
| 24 | ASP55-Main | LEU2-Main | 32.54% |
| 25 | ILE30-Main | ARG26-Main | 31.20% |
| 26 | CYS31-Main | GLY27-Main | 29.20% |
| 27 | LYS1-Main | ASP55-Side | 27.20% |
| 28 | LYS56-Side | ASP94-Side | 23.18% |
| 29 | GLN60-Side | ASP55-Side | 21.64% |
| 30 | LYS43-Main | ALA39-Main | 18.49% |
| 31 | LYS6-Side | GLU57-Side | 18.48% |
| 32 | ALA93-Main | GLY61-Main | 18.44% |
| 33 | GLU50-Main | ILE35-Main | 17.86% |
| 34 | LEU100-Main | GLY23-Main | 15.76% |
| 35 | GLU14-Main | GLU10-Main | 15.16% |
| 36 | GLN88-Side | LYS36-Main | 15.04% |
| 37 | TRP115-Main | LEU111-Main | 13.46% |
| 38 | SER33-Main | LEU29-Main | 12.73% |
| 39 | ARG48-Side | PRO46-Main | 12.18% |
| 40 | TRP44-Main | LYS40-Main | 11.90% |
| 41 | ALA110-Main | GLU14-Side | 11.52% |
| 42 | LYS13-Side | GLU10-Side | 11.45% |
| 43 | LYS56-Side | THR58-Side | 11.33% |
| 44 | GLN84-Side | PHE76-Main | 11.12% |
| 45 | ARG48-Side | CYS120-Side | 11.00% |
| 46 | HSD34-Side | GLY54-Main | 11.00% |
| 47 | GLY59-Main | LYS56-Main | 10.59% |
| 48 | LYS36-Side | GLY54-Main | 10.06% |
| 49 | LYS56-Side | GLU57-Side | 9.78% |
| 50 | LYS13-Side | GLU16-Side | 9.65% |
| 51 | THR97-Side | THR96-Main | 9.55% |
| 52 | LEU103-Main | CYS99-Main | 8.63% |
| 53 | LYS77-Side | GLU72-Side | 8.31% |
| 54 | LEU29-Main | THR25-Main | 8.06% |
| 55 | LYS56-Main | GLN60-Side | 7.81% |
| 56 | ASN18-Side | GLU14-Main | 7.33% |
| 57 | LYS113-Main | SER109-Main | 7.08% |
| 58 | THR96-Side | ALA93-Main | 7.00% |
| 59 | CYS37-Main | GLU50-Side | 6.72% |
| 60 | LYS36-Side | ASP69-Side | 6.72% |
| 61 | ARG48-Side | VAL106-Main | 6.68% |
| 62 | LYS40-Side | LEU79-Main | 6.60% |
| 63 | LYS114-Main | ALA110-Main | 6.53% |
| 64 | LEU116-Main | LEU112-Main | 6.35% |
| 65 | CYS95-Main | CYS92-Main | 6.13% |
| 66 | GLY4-Main | ASP55-Main | 5.96% |
| 67 | THR38-Side | GLN84-Side | 5.75% |
| 68 | TRP52-Main | SER33-Main | 5.72% |
| 69 | LYS36-Side | GLU50-Side | 5.70% |
| 70 | LYS5-Side | GLU53-Side | 5.60% |
| 71 | LYS107-Main | ASN18-Side | 5.51% |
| 72 | ASP55-Main | LYS1-Main | 5.47% |
| 73 | LEU111-Main | GLU14-Side | 5.40% |
| 74 | MET41-Main | THR38-Side | 5.19% |

### Supplementary Figures

**A**

```
GLuc/7D2O       TGKPTENNEDFNIVAVASNFATTDLDADRGKLPGKKLPLEVLKEMEANARKAGCTRGCLI
picALuc         ------------------------------KLPGKKLPLEVLKELEANAQKAGCTRGCLI
                                              **************:****:**********

GLuc/7D2O       CLSHIKCTPKMKKFIPGRCHTYEGDKESAQGGIGEAIVDIPAIPRFKDLEPMEQFIAQVD
picALuc         CLSHIKCTAKMKKWLPGRCESWEGDKETGQGGIGEAIVDIPEIPGFKELAPMEQFIAQVD
                ********.****::****.::*****:.************ ** **:* **********

GLuc/7D2O       LCVDCTTGCLKGLANVQCSDLLKKWLPQRCATFASKIQGQVDKIKGAGGDIEGR
picALuc         LCADCTTGCLKGLANVKCSALLKKWLPSRC------------------------
                **.*************:** *******.**
```

**B**

**Sequence Alignment & Secondary Structure Prediction**

```
ALuc            NHHHHHHHHDIVGVEGKFGTTDLETDLFTIVEDMNVISRDTDVDANRADRGRRGKLPGKK
picALuc         ------------------------------------------------------KLPGKK
Consensus                                                             ******
Sec. Str.                                                             ------

ALuc            LPLEVLKELEANAQKAGCTRGCLICLSHIKCTAKMKKWLPGRCESWEGDKETGQGGIGEA
picALuc         LPLEVLKELEANAQKAGCTRGCLICLSHIKCTAKMKKWLPGRCESWEGDKETGQGGIGEA
Consensus       ************************************************************
Sec. Str.       --HHHHHHHHHHH------HHHHHHH-HHH--HHHHHH----------------------

ALuc            IVDIPEIPGFKELAPMEQFIAQVDLCADCTTGCLKGLANVKCSALLKKWLPSRCAGFADK
picALuc         IVDIPEIPGFKELAPMEQFIAQVDLCADCTTGCLKGLANVKCSALLKKWLPSRC------
Consensus       ******************************************************
Sec. Str.       E-------------HHHHHHHHHH-----HHHHHHHHHHHHHHHHHHHHH----

ALuc            IQAQVDTIKGAGGS
picALuc         --------------
Consensus
Sec. Str.
```

**C**

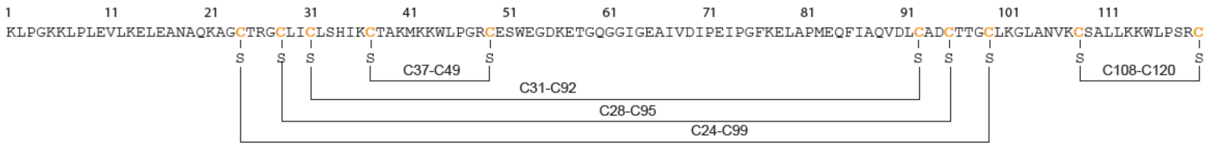


***Supplementary Figure 1.*** *(A) Sequence alignment of GLuc and picALuc. (B) Sequence alignment of picALuc and ALuc. Positively and negatively charged residues are highlighted in blue (light blue, Lys; deep blue, Arg) and red (light red, Asp; deep red, Glu), respectively. Secondary structure prediction is shown in the lower panel. (C) Amino acid sequence of picALuc highlighting all disulfide bridges observed in the structural model of picALuc generate from the NMR structure of Gaussia luciferase (GLuc) [PDB: 7D2O][1].*

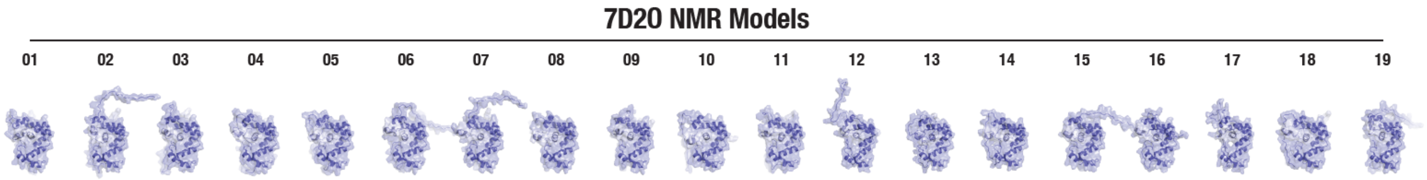


***Supplementary Figure 2.*** *Surface and cartoon representation of GLuc NMR models (conformers; [PDB: 7D2O][1]) revealing flexibility of the N-terminal region of the protein.*

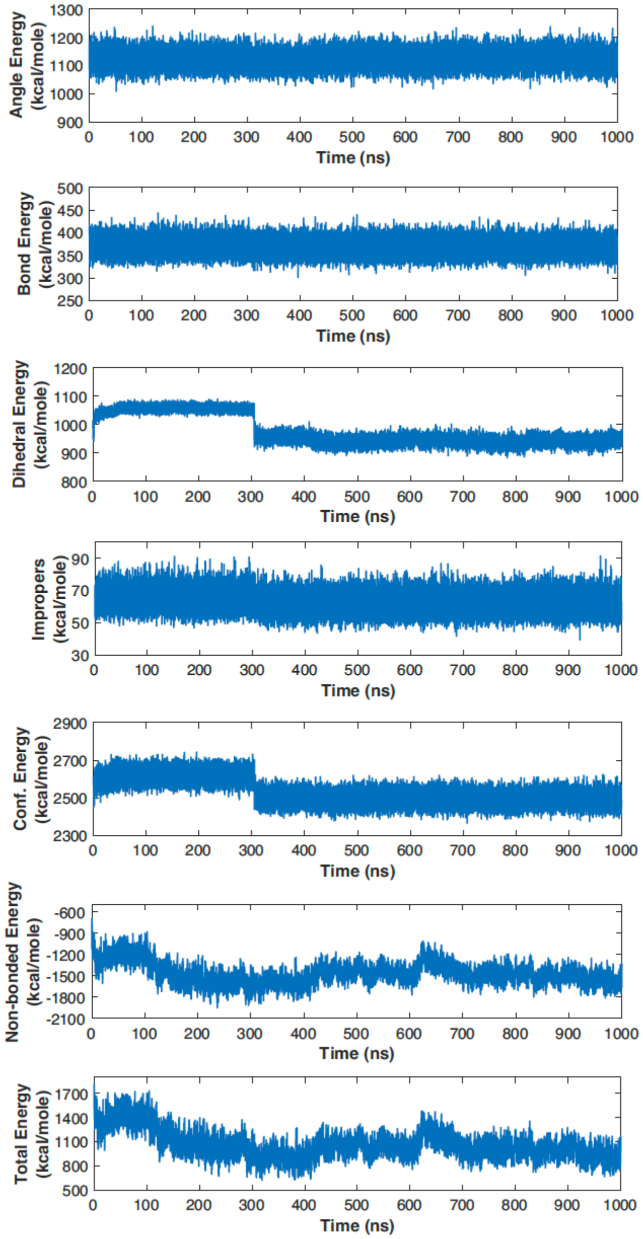


***Supplementary Figure 3.*** *Graphs showing indicated energy term of picALuc over the course of 1 $\mu$s of GaMD simulation. Note the general decrease in total energy of the protein during the initial phases of the simulation.*

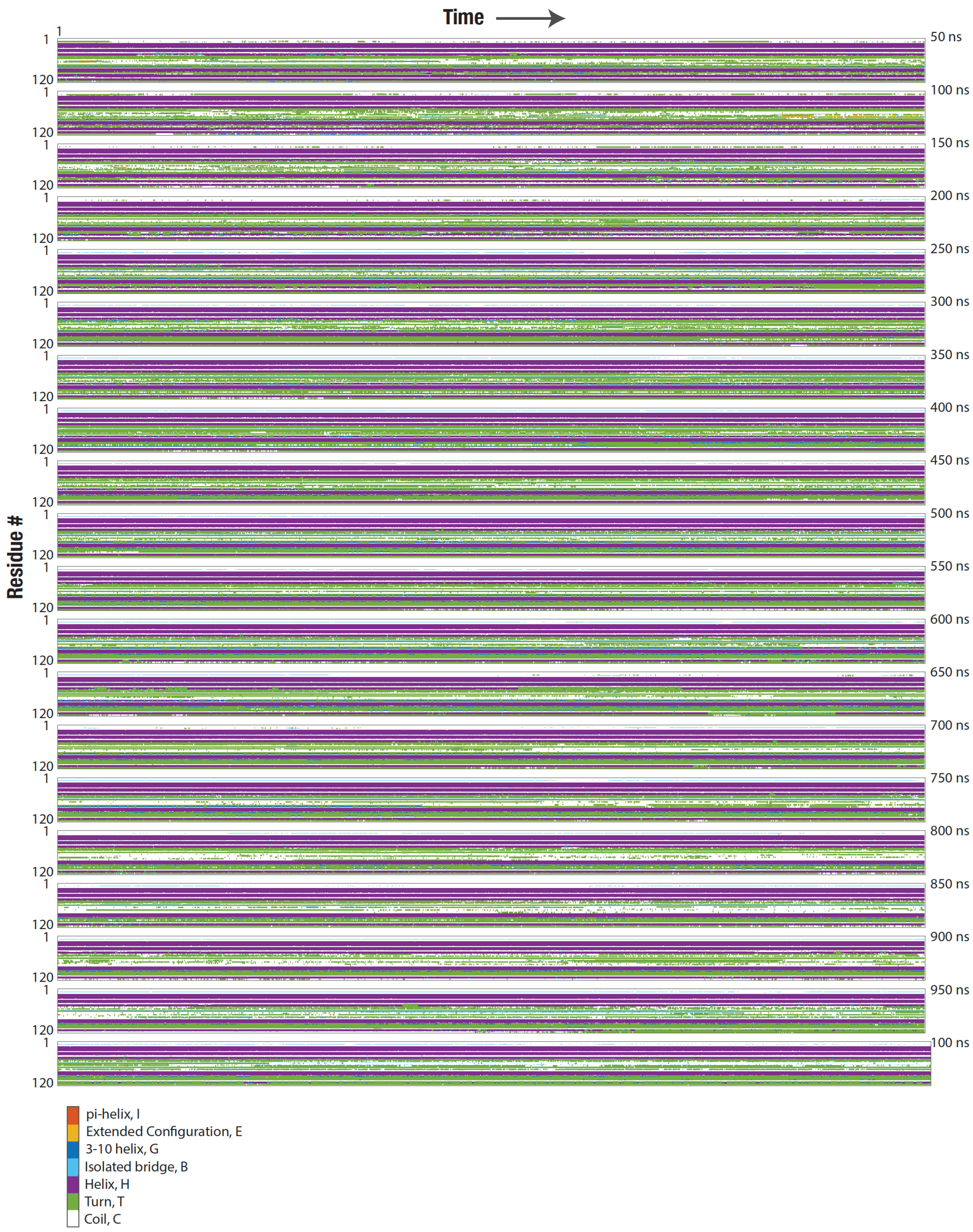


***Supplementary Figure 4.*** *Schematic representation of secondary structure of picALuc over the course of course of 1 μs of GaMD simulation.*

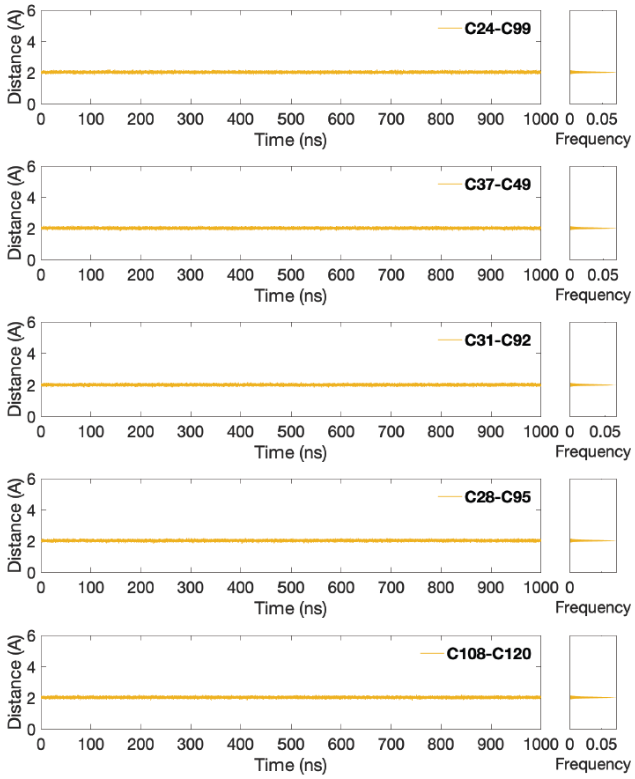


***Supplementary Figure 5.*** *Graphs showing S atom distances between the indicated disulfide forming Cys residues of picALuc over the course of course of 1 $\mu$s of GaMD simulation.*

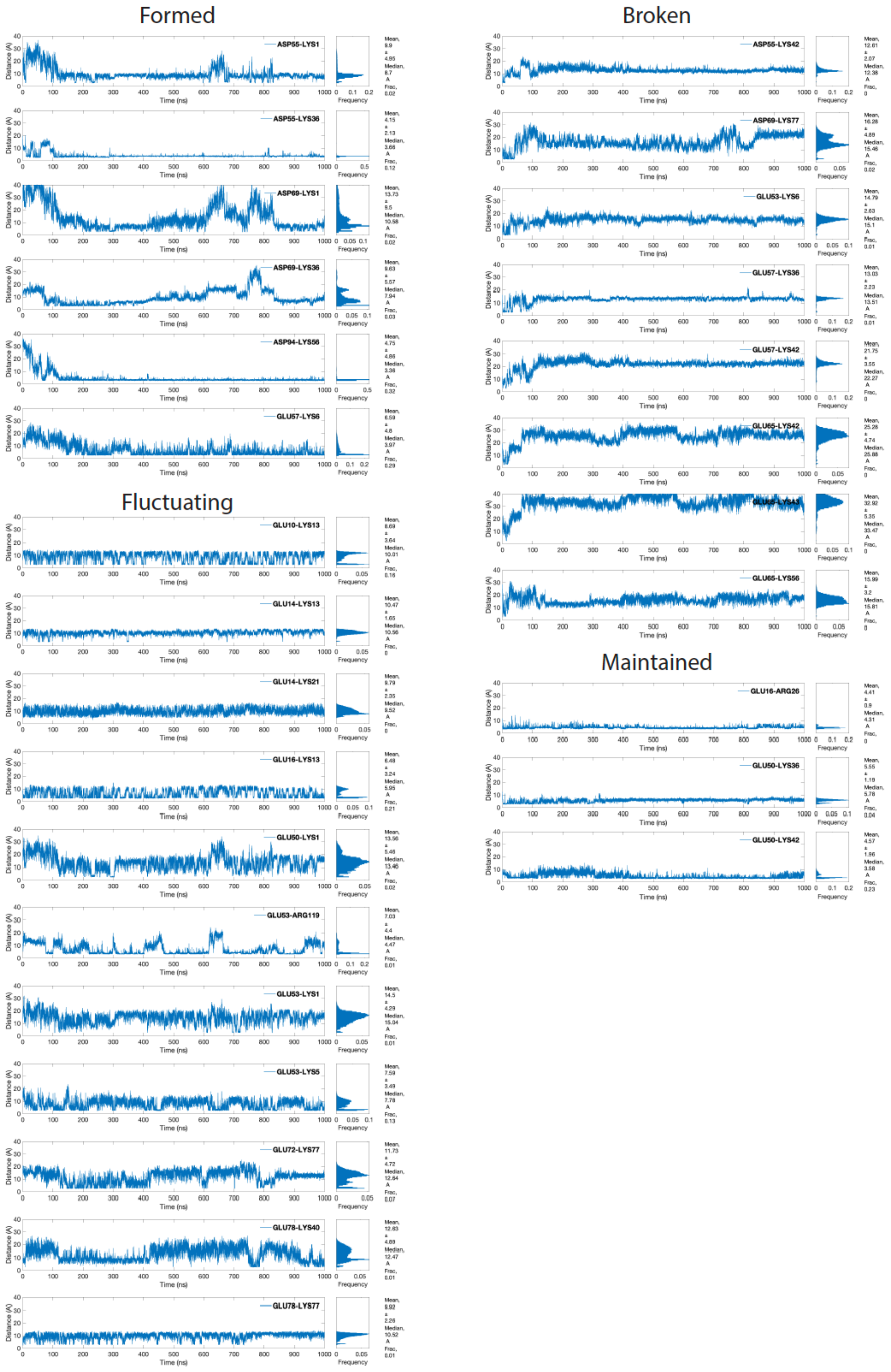


***Supplementary Figure 6.*** *Graphs showing distances between O and N atoms of the salt bridge forming indicated residues over the course of course of 1 μs of GaMD simulation as determined using the Salt Bridges Plugin available in VMD[2]. The interactions have been grouped as those that are formed during the simulation, those that are broken during the simulation, those that fluctuate during the simulation and those that are maintained during the course of the simulation.*

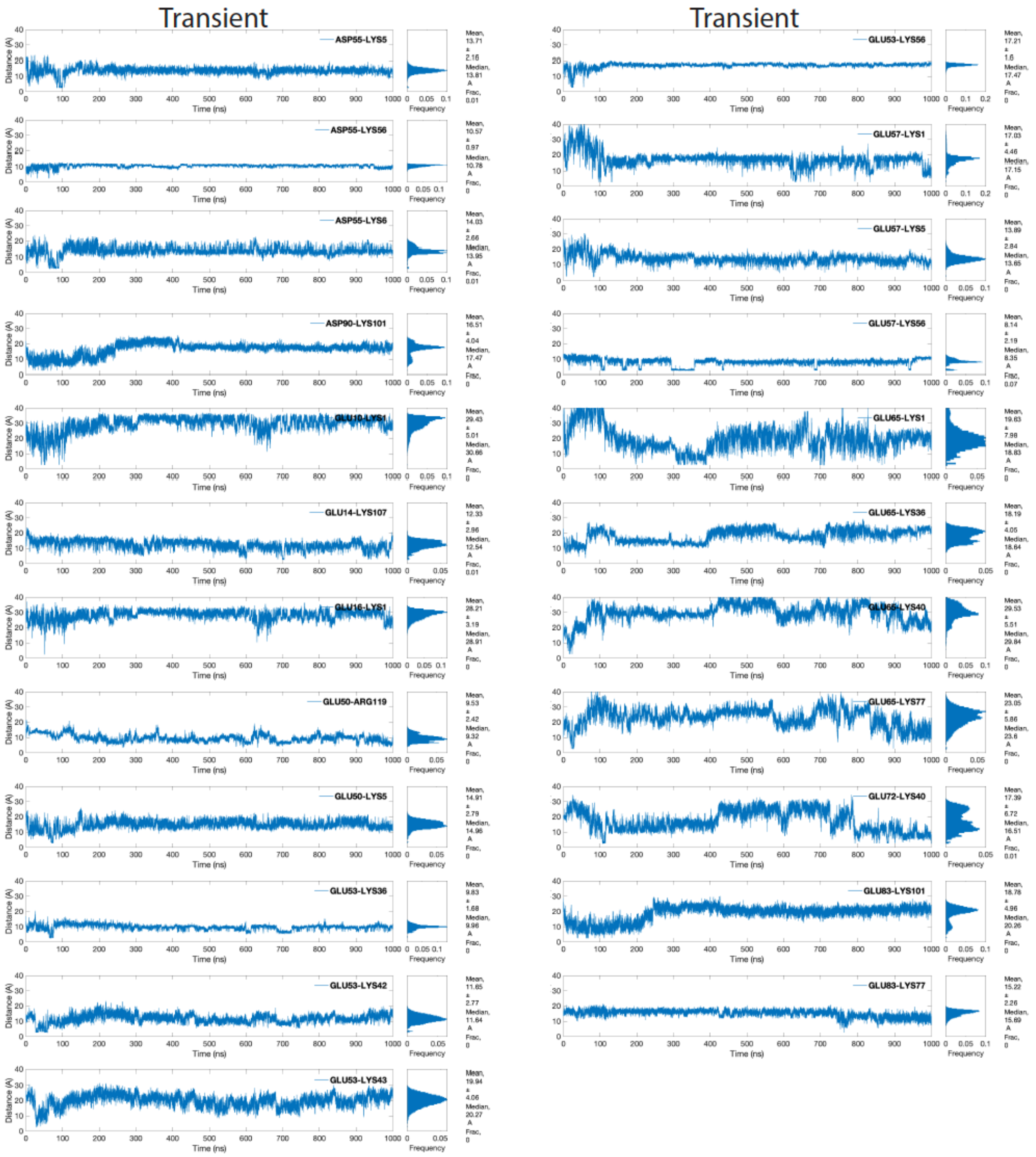


***Supplementary Figure 7.*** *Graphs showing distances between O and N atoms of indicated residues that form transient salt bridges over the course of course of 1 $\mu$s of GaMD simulation.*

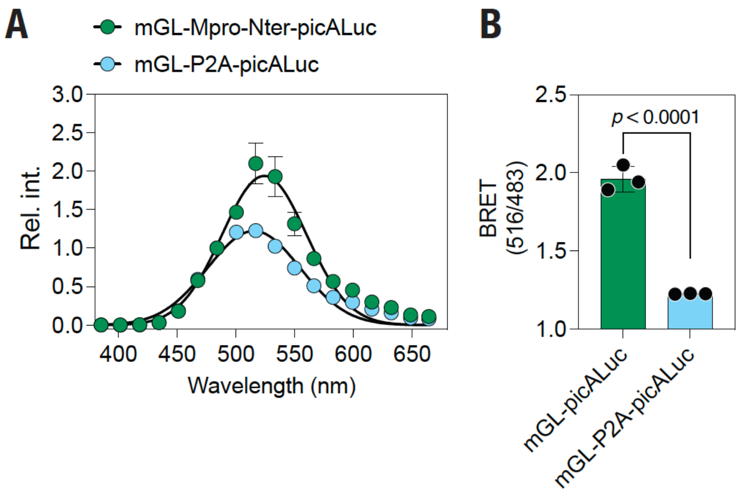

***Supplementary Figure 8.*** *(A) Graph showing bioluminescence spectra of mGL-picALuc and mGL-P2A-picALuc obtained from live cells. (B) Graph showing BRET[3, 4, 5] (ratio of emissions at 516 and 483 nm) of mGL-picALuc obtained from live cells.*

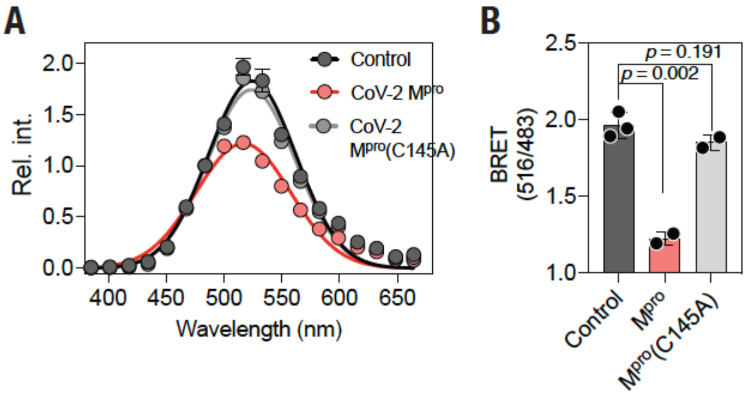


***Supplementary Figure 9.*** *(A) Graph showing bioluminescence spectra of mGL-picALuc obtained from live cells in the absence or presence of either WT or C145A mutant SARS-CoV-2 $M^{pro}$. (B) Graph showing BRET[3, 4, 5] (ratio of emissions at 516 and 483 nm) of mGL-picALuc obtained from live cells in the absence or presence of either WT or C145A mutant SARS-CoV-2 $M^{pro}$.*
